# Behavioural and molecular characterisation of the Dlg2 haploinsufficiency rat model of genetic risk for psychiatric disorder

**DOI:** 10.1101/2021.10.11.463943

**Authors:** Sophie Waldron, Rachel Pass, Simonas Griesius, Jack R. Mellor, Emma S. J. Robinson, Kerrie L. Thomas, Lawrence S. Wilkinson, Trevor Humby, Jeremy Hall, Dominic M. Dwyer

## Abstract

Genetic studies implicate disruption to the *DLG2* gene in copy number variants as increasing risk for schizophrenia, autism spectrum disorders and intellectual disability. To investigate psychiatric endophenotypes associated with *DLG2* haploinsufficiency (and concomitant PSD-93 protein reduction) a novel clinically relevant *Dlg2*^*+/-*^ rat was assessed for abnormalities in anxiety, sensorimotor gating, hedonic reactions, social behaviour, and locomotor response to the N-Methyl-D-aspartic acid receptor antagonist phencyclidine. *Dlg* gene and protein expression were also investigated to assess model validity. Reductions in PSD-93 messenger RNA and protein were observed in the absence of compensation by other related genes or proteins. Behaviourally *Dlg2*^*+/-*^ rats show potentiated locomotor response to phencyclidine, as is typical of psychotic disorder models, in the absence of deficits in the other behavioural phenotypes assessed here. This shows that the behavioural effects of *Dlg2* haploinsufficiency may specifically relate to psychosis vulnerability but are subtle, providing a contrast to the gross deficits in *Dlg2* homozygous models (Winkler, et al., 2018; Yoo et al., 2020a) which do not so specifically model the single chromosome *DLG2* deletion in carriers of risk-associated copy number variants.

## Introduction

The *DLG2* gene locus is linked to multiple psychiatric disorders. Point mutations in promotor regions have been associated with autism ^3^, schizophrenia and intellectual disability ^4^. Copy number variants encompassing complete deletion of *DLG2* result in increased risk for schizophrenia ^5^, autism ^6^, bipolar disorder ^7^ and epilepsy ^8^. Such clinical evidence highlights the potential importance of *DLG2* in the psychopathologies common to a broad range of disorders. *DLG2* encodes the postsynaptic scaffold protein PSD-93 (also known as Chapsyn-110) in the membrane associated guanylate kinase (MAGUK) family. These proteins are responsible for anchoring and organising the numerous protein complexes required for development and plasticity at the synapse, in particular the NMDA receptor ^9–12^, AMPA receptor ^13^, potassium ion channels ^10,14,15^ and neuroligin 1-3 ^9^.

Previous investigation into specific behavioural endophenotypes driven by *Dlg2* disruption using *Dlg2*^*-/-*^ mice have shown abnormal social behaviours including impaired social preference ^16^, increased repetitive behaviours and hypoactivity in response to novelty ^16,17^. Deficits in cognitive flexibility and attention have also been shown ^18^, aligning with similar deficits in human carriers of mutations to the *DLG2* coding region. Other psychiatric endophenotypes are yet to be tested, including faulty sensorimotor gating, anhedonia, and locomotor response to pharmacological challenge.

A limitation of the previous studies of the molecular and behavioural consequences of alteration to *Dlg2* is that they have thus far focused on homozygous knockouts ^16,17^ rather than the more clinically relevant heterozygous knockouts. Heterozygous models capture the single chromosome deletion of *DLG2* in a CNV ^5,19–21^ or single chromosome loss-of-function mutations ^3,4^. This work presents the first molecular and behavioural characterisation of a rat model generated using CRISPR/Cas9 gene editing technology which contains only one copy of the *Dlg2* gene.

Here we assess whether the expected biological changes (reduced *Dlg2* mRNA and protein expression) occurred in the heterozygous (+/-) model, and whether there is evidence of compensation for possible *Dlg2* reduction from other MAGUK family members or related proteins, as has been shown in cortical neurons with complete PSD93 knockdown ^22^. Behavioural consequences of *Dlg2* haploinsufficiency was assessed by comparing *Dlg2*^*+/-*^ rats and wild-type littermates on a battery of psychiatric-relevant translational tests. These included tests of anxiety, social behaviour, and anhedonia, key behavioural domains disrupted across disorders *Dlg2* is implicated in. Regulating unwanted or unnecessary sensory inputs was assessed using sensorimotor gating, deficiencies of which characterise patients with schizophrenia ^23,24^ and autism ^25^, in addition to rodent models of these conditions ^26^. Hyperlocomotion in response to PCP (an NMDAR-antagonist) was also assessed as acute administration of PCP produces a transient psychosis-like phenotype ^27,28^ which may be exaggerated in a rodent model with potential NMDAR alteration of function ^29^ and an underlying genetic propensity towards psychosis.

*Dlg2*^*+/-*^ rats demonstrated selective reductions in mRNA and protein expression that was not compensated for by increases in the remainder of the *Dlg* family. There was also an absence of gross behavioural deficits related to anxiety, hedonic reactions, social behaviour, and sensorimotor gating in the model; however, *Dlg2*^*+/-*^ rats demonstrated a potentiated hyperlocomotion phenotype in response to PCP administration compared to wild-types. Together, these confirm the biological validity of the current model selectively relevant to reduced expression of *Dlg2* and a possible concomitant alteration of NMDAR function ^29^ and furthermore suggests that clinically relevant reductions to *Dlg2* expression will have more subtle effects than homozygous knockout models.

## Materials and Methods

### Animals

*Dlg2* heterozygous rats were generated on a Long Evans Hooded background by Horizon Discovery (Pennsylvania, USA) using CRISPR/Cas9 gene editing technology. Successful founders generated by Horizon Discovery has a 7bp deletion (782933-782939 in the genomic sequence) in exon 5 which caused a frame shift and generation of an early stop codon in exon 6. Confirmation of successful non-homologous end joining activity was assessed by PCR and sequenced by Horizon Discovery, UK. Selected heterozygous founders were send to Charles River (Margate, UK) and bred to produce experimental colonies by breeding male heterozygous rats were bred with female wild-types resulting in a Mendelian distribution of wild-type and heterozygous pups. A more detailed description of the generation of this rat line can be found in Supplement 1 of Griesius et al. ^29^

Animals were housed in groups from two to four in standard cages (l × w × h: 50cm × 32cm × 21cm) in rooms with a temperature between 19-23°C maintained on a 12h light-dark cycle. Cages had sawdust and paper nesting and environmental enrichment (wooden chews and cardboard tubes). Food and water were given ad-lib while conducting all tasks except the lick microstructure assessment where rats were maintained at 85-95% of their free-feeding weight by giving them restricted access to food at the end of each day. Research was conducted in accordance with the Home Office regulations under the Animal (Scientific Procedures) Act 1986 Amendment Regulations (SI 2012/3039) under the authority of PPL 303243 or PPL 303135. Five cohorts of animals were used across the studies here. Cohort 1 – 20 male (8 wild-type 12 *Dlg2*^*+/*-^), 25 female (16 wild-type 9 *Dlg2*^*+/*-^) for assays elevated plus maze, open field and sensorimotor gating in that order; Cohort 2 – 48 male (28 wild-type 20 *Dlg2*^*+/*-^) for lick microstructure assessment; Cohort 3 – 33 male (14 wild-type 19 *Dlg2*^*+/*-^), 25 female (12 wild-type 13 *Dlg2*^*+/*-^) for assays social preference and PCP hyperlocomotion in that order; Cohort 4 – 24 male (12 wild-type 12 *Dlg2*^*+/*-^) for Western blot and Cohort 5 – 16 male (8 wild-type 8 *Dlg2*^*+/*-^) for qPCR.

### Tissue extraction

Rodents used for qPCR and Western blot were culled by inhalation of slowly rising C0_2_ concentration for eight minutes (administered by Home Cage Culling Chamber, Clinipath Equipment Ltd, UK) two weeks after completion of behavioural experiments not reported here at 2-4 months old. Brains were extracted from the skull and gross dissected by partitioning the cerebellum and rostral-most part of cortex (prefrontal cortex). The hippocampus and posterior cortex were then dissected. Extracted brain regions were flash frozen on dry ice and stored at −80°C prior to use.

### Quantitative polymerase chain reaction (qPCR)

Using the QIAGEN RNeasy kit, RNA was isolated from prefrontal cortex, hippocampal and cerebellar tissue from individual animals (Cohort 5). Samples were DNAase treated using TURBO DNA-free™ Kit (Ambion Life Technologies), following the recommended protocol. cDNA synthesis was performed using the RNA to cDNA EcoDry™ Premix (Random Hexamers) synthesis tubes (Clontech), heated at 42°C for 75 minutes, followed by 80°C for 15 minutes. qPCR was conducted with SensiMix SYBR Green (Bioline) on the StepOne Plus (Life Technologies; 1 cycle 95°C, 10 mins; 45 cycles of 95°C, 15 secs and 60°C, 1 min; with melt curves conducted 55°C, 1 min; 95°C for 15 secs). All qPCR samples were run in triplicate and the outcome was calculated using 2^-ΔΔCt^ method, normalised to UBC and SDHA housekeeping genes. Primer sequences are given in Table 1.

**Table 1:**
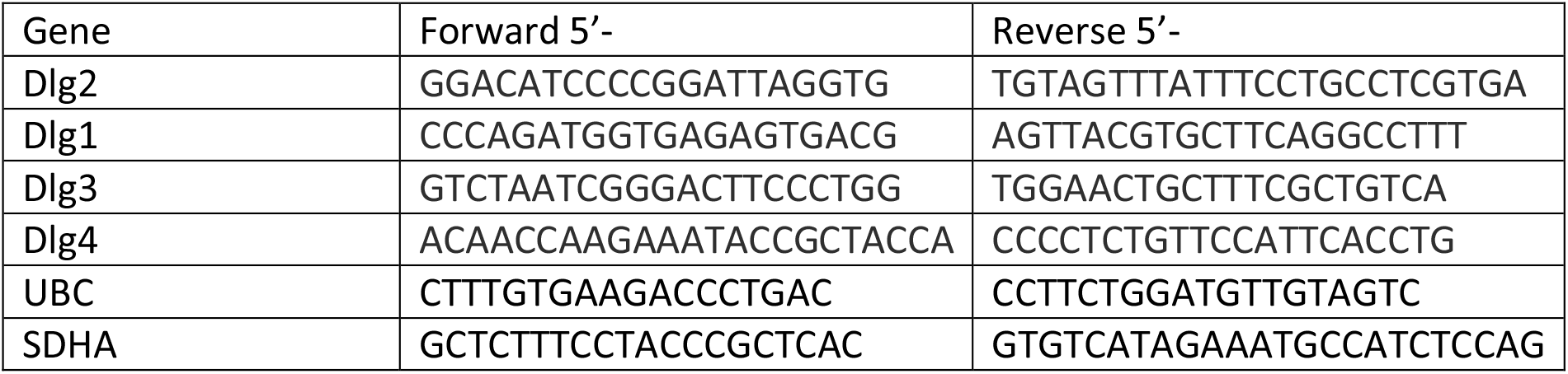
Primer sequences used in qPCR.

### Western blot

Western blot analysis was conducted on hippocampal, prefrontal cortex, posterior cortex and cerebellar tissue from WT (n=12) and HET (n=12) animals. Each tissue sample was lysed in Syn-PER lysis and extraction buffer (Thermo Fisher, UK) with mini protease inhibitor cocktail (Roche Diagnostics) and phosphatase inhibitor (Cell signalling, UK) according to description from manufacturer. After using BCA Assay kit to measure the total amount of protein in each sample, electrophoresis and blotting were carried out.

Gels (4–12% NuPAGE Bis-Tris Midi, 45 well) were loaded with 40ug of protein was loaded per well. Samples were added to Laemmli buffer at a 1:1 ratio and this mixture heated at 96 °C for 5 minutes to denature protein-protein interactions and facilitate antibody bindings. Samples were arranged so that brain regions and genotypes were counterbalanced across gels with a WT standard used on each gel for comparison. Gels were run at room temperature in NuPAGE™ Running Buffer (Invitrogen, UK) at 85V for 20 mins and then for a further hour at 115V. Protein was then transferred to 0.45 um pore size nitrocellulose membrane (Invitrogen, UK) at 85 V for 2 hours 15 minutes at an ambient temperature of 4 °C in NuPAGE™ Transfer Buffer (Invitrogen, UK) containing 10% 2-propanol (ThermoFisher Scientific, UK). Membranes containing transferred protein were washed in Tris-Buffered Saline (20mM Tris, 150mM NaCl, pH 7.6) with 0.1% Tween 20 (TBST) before blocking in 5% milk for one hour at room temperature with gentle rocking.

Primary antibodies were diluted to appropriate concentrations in 5% milk and incubated with the membrane overnight at 4 °C. These included rabbit anti-PSD93 (1:1000, Cell Signalling Technology, USA), mouse anti-NR1 (1:1000, Merck Millipore, UK), rabbit anti-PSD95 (1:2000, Abcam, UK) and mouse anti-GAPDH (1:5000, Abcam, UK). Membranes were then subject to 3 × 10 minute TBST washes before incubation with the appropriate fluorescent IRDye 680RD secondary antibodies at 1:15,000 dilution in 5% milk at room temperature. After another series of TBST washes membranes were imaged on Odyssey CLx Imaging System (Li-COR, Germany). Densiometric analysis of bands was performed using ImageLab 6.0 (https://imagej.nih.gov/ij/). The densities (with background subtracted) of the protein of interest were divided by the loading control densities for each sample to provide normalised values. Densities were then averaged by group. The hippocampal protein expression outlined here is also reported in Griesius et al ^29^ Supplementary Information.

### Elevated plus maze

The elevated plus maze (EPM) consisted of two open arms (45 cm long × 10 cm wide), two closed arms (45 cm long × 10cm wide × 30cm high) and a middle (10 × 10 cm) compartment forming the shape of a plus sign, elevated 50cm above the ground. The room was dimly lit, with the light level in the open arms 26 lux, and the light level in the closed arms 15.3 lux. Rats were habituated to the testing room for at least one hour before individual testing. Each rat was placed in the middle compartment with its head facing an open arm and allowed to freely explore the apparatus for five minutes. Between tests the arena was cleaned with 70% ethanol. Each test was recorded by a camera mounted 120 cm above the maze and MP4 videos subsequently analysed for movement across the maze by Ethovision software (Version XT V 13, Noldus Information Technologies, Netherlands) (frame rat of 7/s). For the EPM, virtual zones for each of the open and closed arms and the middle were created and total time spent with all four paws in each region, plus movement and velocity across the whole maze were analysed. Head-dips, stretch attend postures and grooming were manually scored. A head dip is defined when a rat is on an open arm and peers over the platform edge such that it’s head is fully off the platform and a stretch attend posture (SAP) is scored when the body of the rat is close to the floor and it’s rear legs are in the closed arm whilst it investigates the open arms or middle section. Anxiety on the EPM is associated with more stretch attend postures, fewer head dips and increased defecation in addition to reduced open arm exploration ^30^. Faecal boli were also counted for each animal after each run, and for female rodents vaginal cytology was performed after testing on the maze to determine oestrus stage

### Open field test

The open field apparatus was a 100 × 100 cm square wooden arena with 30cm high walls, painted black. Light levels were 25 lux at the centre of the maze, and 11.8 lux in the corners of the maze. Rats were habituated to the test room for at least one hr, and were then placed, individually, in the arena adjacent to the middle of the south wall, with their head facing the wall. Between tests the arena was cleaned with 70% ethanol. Animal movements were recorded by a camera mounted 200 cm, above the arena. Ethovision with virtual zones dividing the arena into a central region centre (25cm^2^ located 25cm from each other wall) and outer region (75cm^2^, within 25cm of each wall was used to analyse the recordings. Locomotor activity and thigmotaxis were assessed during a 10-minute session, in terms of the amount of time rodents spent in the central and outer regions, the distance moved (cm) and velocity of movement(cm/s) in the entire apparatus. Faecal boli were also counted for every rodent and for female rodents vaginal cytology was performed after testing on the maze to determine oestrus stage.

### Acoustic startle response (ASR) and pre-pulse inhibition (PPI)

ASR and PPI were assessed using a pair of R-Lab™ Startle Response System chambers (San Diego Instruments, San Diego, USA). Each sound proofed chamber was equipped with a Perspex enclosure (10 cm diameter), with doors at either end, into which a rat was placed. This tube was located on a Perspex plinth, directly of a Piezoelectric sensor which register flexion and converted this to an electrical signal (ASR) that was monitored by a computer equipped with SR-LAB startle software (San Diego Instruments, San Diego, USA). In the roof of the chamber, above the centre of the enclosure was a loudspeaker through which background noise (65db) and trial stimuli were presented.

Methods were conducted as in Geyer, Wilkinson, Humby, and Robbins ^31^. A session (30 min, 91 trials) consisted of a five-minute period of habituation (at background noise) followed by two blocks of trials. Blocks 1 and 2 consisted of 13 pulse-alone startle trials (40 ms duration, at either 120 or 105db) and 15 prepulse trials, 5 trials each at either 4, 8 or 16db above background noise levels, intermixed on a pseudorandom schedule. A prepulse trial consisted of a stimulus (40 ms duration) followed by a startle stimulus, as above, (100 ms onset to onset delay). The third block of 18 trials of pulse-alone startle trials, 3 of each at 70, 80, 90, 100, 110 and 120 db, pseudorandomly presented. The ASR to the first three pulse-alone trials at 120 dB and 105 dB in blocks 1 and 2 were averaged and analysed as an index of emotional reactivity, mean ASR was calculated as the average of the remaining 10 pulse-alone trials/block and response to prepulse trials at each intensity/block were also averaged. PPI was calculated as the proportional difference between mean ASR for pulse-alone trials and mean prepulse response at each prepulse intensity. ASR for trials in Block 2 were meaned together at each intensity used. Background intensities 105 dB and 120 dB were used to mitigate against potential floor and ceiling effects. However, with no evidence of such effects, and the pattern or results observed at both intensities, only the findings for a background intensity of 120 dB are reported here. All ASR values were weight adjusted before analysis. As previously, rats were habituated to a room for at least 60 min before testing, and then transferred to the test room and individually placed into each enclosure within a test chamber. Enclosures were cleaned 70% ethanol between subjects.

### Lick microstructure assessment

Rats were trained and tested in 16 custom made drinking chambers (Med Associated Inc., St Albans, USA). These were 30 × 13 × 13 cm (L × W × H), with steel grid flooring and white plastic walls. Sucrose was accessible through drinking spouts attached to 50 ml cylinders, which could be lowered through left or right apertures in the front wall of the chamber by hand. A contact sensitive lickometer registered the licks made by rats to the nearest 0.01s once the bottle was available, and MED-PC Software (Med Associates, Inc) recorded the data. Rats were trained across five consecutive days for 10 minutes each day to drink 8% sucrose solution from the spouts. During the first session the spout was left to protrude into the cage to encourage drinking, but after this the spout stopped just beyond the opening in the cage to minimise accidental contact. Once all rats were consistently drinking, the test phase began. During test rats drunk 4% and 16% sucrose solutions for four days each in an order counterbalanced for genotype. Rats were allocated to cages in alternating wild-type/ *Dlg2*^*+/-*^ order and the same drinking cage was used for each animal across the experiment. Half of the rats received four days of 16% followed by four days of 4% and the other half the reverse to implement the counterbalance. The amount of fluid consumed by each rat was measured by weighing the drinking bottle before and after each session. Solutions were made up daily on a weight/weight basis.

Mean consumption of sucrose (g) and mean lick cluster size for each rat were extracted from the record of licks for analysis. A cluster was defined as a set of licks, each separated by an interlick interval of no more than 0.5 s. This criterion was used by Davis and his co-workers who pioneered this technique ^32,33^ and in many previous studies employing this assay to assess analogues of anhedonia ^34–37^.

### Social preference test

This test utilised the same arena as the open field with two wire mesh chambers (22 cm diameter) weighed down with 2kg weights placed diagonally in opposite corners of the arena. The distance between side walls and the chambers was 18 cm, and the distance between them diagonally always 35 cm. Light was 24.8 lux. Rats were placed in the experimental room where the arena was separated from them by a curtain for 1h prior to testing commencing. Each individual rat was given 10 minutes to explore the arena with empty chambers then removed to a separate holding cage for five minutes, before being placed back in the arena for the 10-minute social preference test. In the social preference test the rat was presented with test one chamber contained an unknown conspecific (stranger rat) and the other chamber and unknown object. For each test animal a same-sex wild-type rat that had no prior contact with the test rat was used as the stranger. All stranger rats were habituated to the chambers for 10 minutes prior to testing. For analysis raw exploration times of the chambers (rodents directing their nose at the chamber at a distance of < 2 cm) were used in addition to d2 discrimination ratio (equation 1). Discrimination ratio gives a readout of the difference in exploration time between the two stimuli without the confound of overall tendency to explore for long or short durations.

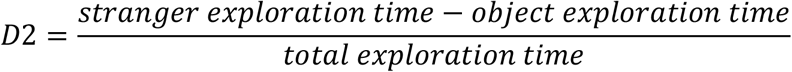

### Phencyclidine (PCP) induced locomotion

To examine PCP-induced changes to locomotor activity rats were placed in 58 × 45 × 60 cm (l x w x h) Perspex boxes and recorded with a camera placed 200 cm above the boxes. Four boxes were used simultaneously but 60 cm high barrier walls prevented rats from interacting with each other. Rats were placed in the boxes for a 30-minute habituation period before being injected subcutaneously with 5 mg/kg dose of PCP hydrochloride (in 0.9% (w/w) saline, Sigma-Aldrich, UK) and returned to the same box for a further 90 minutes. Ethovision software was used to analyse movement and velocity for each rat throughout the habituation and post-drug phases in 10 min blocks.

### Statistical analysis

All statistical analysis was performed using JASP Version 0.14.1 (JASP Team (2020). For traditional null-hypothesis significance testing p < 0.05 was considered statistically significant. For ANOVA analysis where Maunchly’s test indicated sphericity was violated Greenhouse-Geisser corrected values are reported. Because interactions that included sex were non-significant in all analyses that included sex as a factor, the effects of sex as a variable are not reported here but these can be seen in Supplement 1.

Traditional null-hypothesis significance testing only assesses how unlikely the observed data is given the assumption of the null hypothesis, and thus p > 0.05 does not distinguish evidence for the null hypothesis from data insensitivity ^38^. In contrast, Bayesian tests calculate the relative probabilities of the null and alternative hypotheses, and thus allow assessment of whether the evidence is in favour of either hypothesis. In this body of work Bayesian statistics have been applied where traditional null-hypothesis significance testing shows a non-significant result for a key effect or interaction where a null result is potentially theoretically informative (in particular, evidence for a lack of a difference across genotypes).

Bayes factors relate to the ratio of probability for the observed data under a model based on the null hypothesis compared with a model based on some specified alternative. When represented as BF01 Bayes factors vary between 0 and infinity, where 1 indicates that the data do not favour either model more than the other, values greater than 1 indicate increasing evidence for the null over the alternative hypothesis and values less than 1 increasing evidence for the alternative over the null hypothesis. When using Bayes factors to decide whether there is substantial evidence for the null over the alternative, the following conventions suggested by Jeffreys et al. ^39^ can be followed: a Bayes factor between 1 and 3 gives weak or anecdotal support to the null, a factor between 3 and 10 represents some supporting evidence, while a factor more than 10 indicates strong evidence for the null.

Bayes factors were calculated for factorial ANOVAs in the way described by Rouder, Morey, Speckman, and Province ^40^ and Rouder, Morey, Verhagen, Swagman, and Wagenmakers ^41^ and were implemented using JASP 0.14.1 and the default prior scale for fixed and random effects and reported as the analysis of effects – this gives a BF_exclusion_ which is equivalent to BF01 when averaging across models including the factor or interaction of interest. Bayes factors for t-tests were calculated as described by Rouder, Speckman, Sun, Morey & Iverson ^42^ and implemented using JASP 0.14.1 with the default settings for the Cauchy prior distribution on effect size under the alternative hypothesis.

## Results

### Protein and mRNA expression

mRNA expression for the *Dlgs* is shown in Figure 1 with summary values shown for the prefrontal cortex and hippocampus for *Dlg2* (A-B), *Dlg1* (C-D), *Dlg3* (E-F) and *Dlg4* (G-H). ΔCt was analysed by repeated measures ANOVA and Bayesian repeated measures ANOVA with repeated measures factors of brain region (prefrontal cortex, hippocampus) and between-subjects factors of genotype. *Dlg2* expression varied with genotype (genotype main effect: *F*(1, 13) = 29.367, *p* < 0.001, *n*^*2*^_*p*_ = 0.693) but did not vary with brain region (brain region main effect: *F*(1, 13) = 0.690, *p* = 0.421, *n*^*2*^_*p*_ = 0.050; BF_exclusion_ = 2.049 and interaction: *F*(1, 13) = 0.031, *p* = 0.863, *n*^*2*^_*p*_ = 0.002; BF_exclusion_ = 1.691).

**Figure 1:**
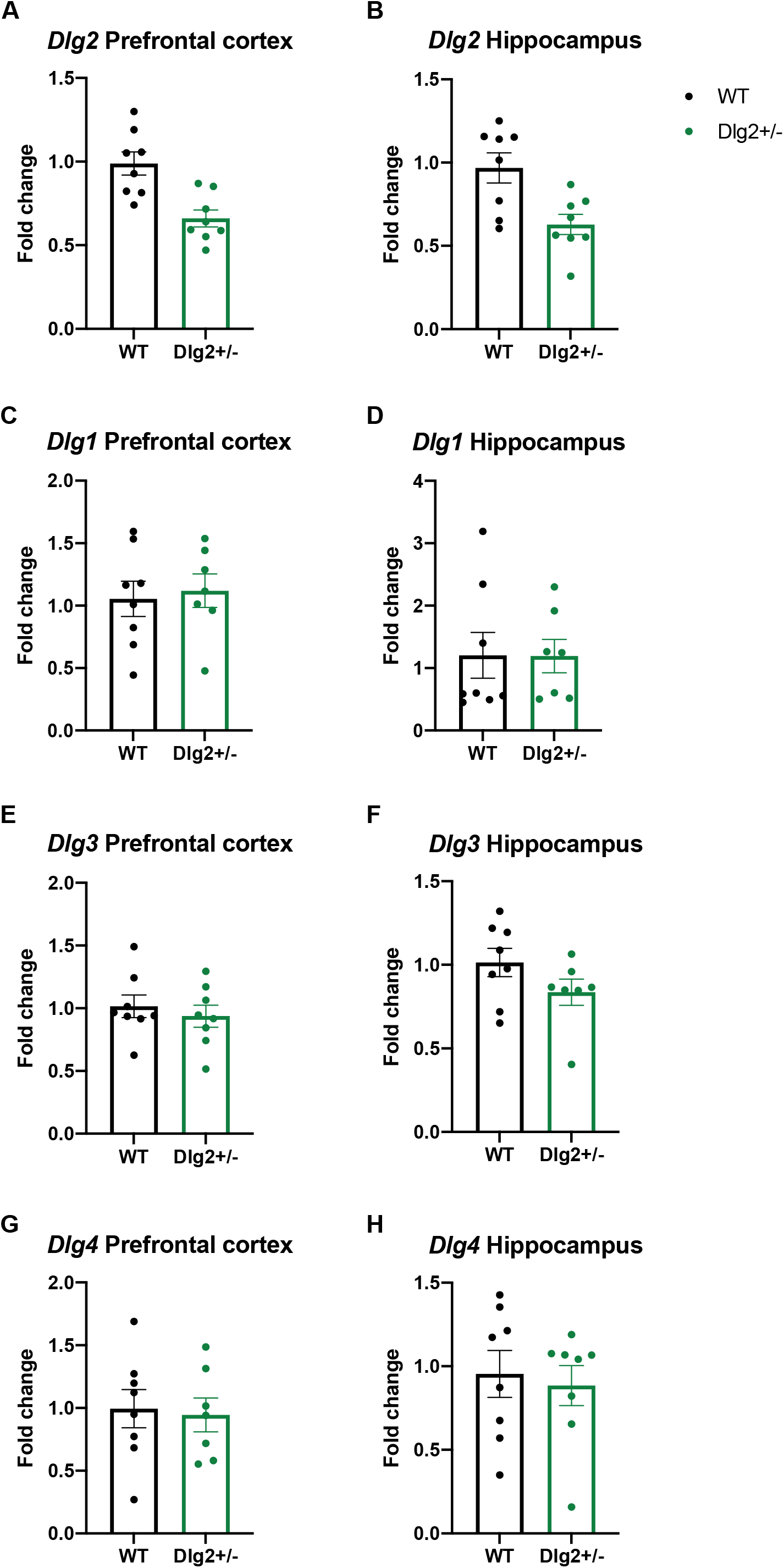
mRNA expression of *Dlg1-4* in *Dlg2*^*+/-*^ and wild-type rats. Data is shown as mean ± SEM fold change plotted plus individual data points for *Dlg2* PFC (A), *Dlg2* hippocampus (B), *Dlg1* PFC (C), *Dlg1* hippocampus (D), *Dlg3* PFC (E), *Dlg3* hippocampus (F), *Dlg4* PFC (G) and *Dlg4* hippocampus (H).

The expression of *Dlg1* differed with brain region (*F*(1, 12) = 7.017, *p* = 0.021, *n*^*2*^_*p*_ = 0.369) yet there were no genotype main effects (*F*(1, 12) = 0.498, *p* = 0.494, *n*^*2*^_*p*_ = 0.040; BF_exclusion_ = 2.307) or interactions (*F*(1, 12) = 0.048, *p* = 0.831, *n*^*2*^_*p*_ = 0.004; BF_exclusion_ = 1.650). This was much the same for *Dlg3*: brain region main effect (*F*(1, 12) = 11.150, *p* = 0.006, *n*^*2*^_*p*_ = 0.482), with non-significant results for genotype (*F*(1, 12) = 2.526, *p* = 0.138, *n*^*2*^_*p*_ = 0.174; BF_exclusion_ = 1.307) and brain region × genotype (*F*(1, 12) = 0.528, *p* = 0.481, *n*^*2*^_*p*_ = 0.042; BF_exclusion_ = 1.003). For *Dlg4* there were no main effects of brain region (*F*(1, 12) = 1.397, *p* = 0.260, *n*^*2*^_*p*_ = 0.104; BF_exclusion_ = 1.726), genotype (*F*(1, 12) = 0.155, *p* = 0.701, *n*^*2*^_*p*_ = 0.013; BF_exclusion_ = 2.786) and no brain region × genotype interaction (*F*(1, 12) = 0.212, *p* = 0.653, *n*^*2*^_*p*_ = 0.017; BF_exclusion_ = 3.852). This indicates that at the mRNA level there is no evidence of compensation for *Dlg2* decreases by changes in expression of other *Dlgs*.

Of the three proteins analysed only PSD-93 showed consistent decreases across all four brain regions in the *Dlg2*^*+/-*^ rats compared to wild-types (Figure 2). Integrated densities were analysed using repeated measures ANOVA with within-subjects factor of brain region (prefrontal cortex, posterior cortex, hippocampus, cerebellum – apart from NR1 where expression in cerebellum was negligible in all cases and thus this region was omitted from the analysis) and between-subjects factor of genotype. Example blots can be seen in Supplementary Figure S1. Repeated measures ANOVA analysis for PSD-93 showed a significant main effect of genotype (*F*(1, 22) = 13.680, *p* = 0.001, *n*^*2*^_*p*_ = 0.383) demonstrating the success of the heterozygous gene knockout on reducing PSD93 protein levels. There was also a significant main effect of brain region (*F*(1.018, 22.400) = 20.893, *p* < 0.001, *n*^*2*^_*p*_ =0.487) and genotype × brain region interaction (*F*(1.018, 22.400) = 5.166, *p* = 0.032, *n*^*2*^_*p*_ = 0.190). The significant genotype × brain region interaction was followed up with independent samples t-tests. PSD-93 was more abundant in the PFC of wild-types than *Dlg2*^*+/-*^ rats (*t*(22) = 6.057, *p* < 0.001, *d* = 2.473), likewise in the hippocampus (*t*(22) = 5.378, *p* < 0.001, *d* = 2.195), posterior cortex (*t*(22) = 4.032, *p* < 0.001, *d* = 1.646) and cerebellum (*t*(22) = 2.702, *p* = 0.013, *d* = 1.103).

**Figure 2:**
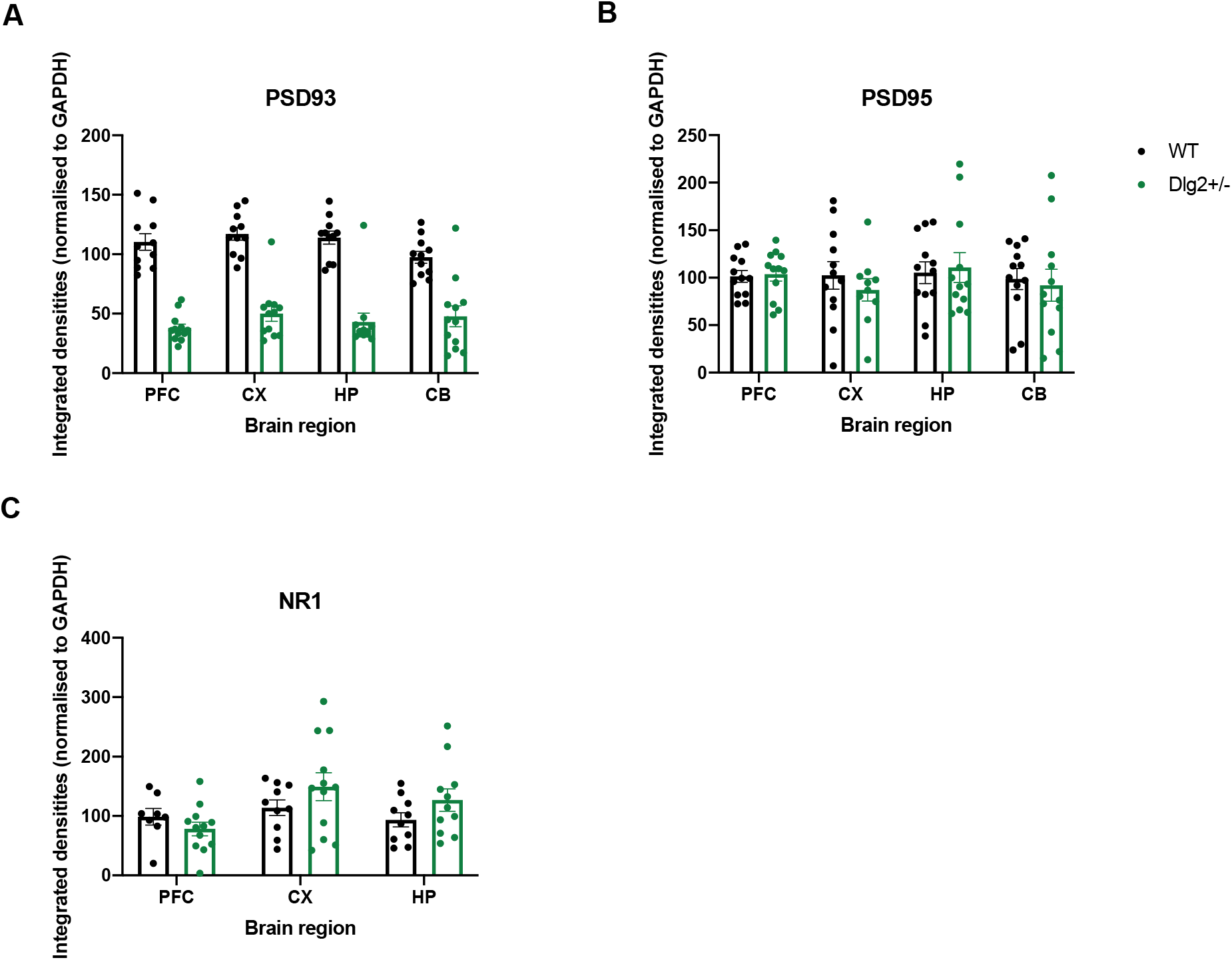
Expression of proteins PSD-93 (A), PSD-95 (B) and NR1 NMDA receptor subunit (C) in *Dlg2*^*+/-*^ and wild-type rats. These were assessed across four brain regions: prefrontal cortex (PFC), posterior cortex (CX), hippocampus (HP) and cerebellum (CB). Cerebellar NR1 expression was too low for analysis thus is not reported. Data is shown as mean ± SEM integrated density plotted plus individual data points.

Repeated measures ANOVA analysis of PSD-95 levels showed no main effect of genotype with Bayes factor inconclusive (*F*(1, 22) = 3.805, *p* = 0.064, *n*^*2*^_*p*_ = 0.147; BF_exclusion_ = 2.187), no main effect of brain region (*F*(1.722, 37.884) = 0.175, *p* = 0.808, *n*^*2*^ = 0.008; BF_exclusion_ = 18.288) and no genotype × brain region interaction with Bayes factors providing evidence for the null (*F*(1.722, 37.884) = 1.140, *p* = 0.324, *n*^*2*^_*p*_ = 0.049; BF_exclusion_ = 23.396).

Similarly for analysis of NR1 NMDA receptor subunit levels there was no main effect of genotype (*F*(1, 17) = 0.262, *p* = 0.616, *n*^*2*^_*p*_ = 0.015; BF_exclusion_ = 3.752), brain region (*F*(1.148, 19.514) = 0.884, *p* = 0.373, *n*^*2*^_*p*_ = 0.049; BF_exclusion_ = 4.026) or genotype × brain region interaction (*F*(1.148, 19.514) = 0.902, *p* = 0.368, *n*^*2*^_*p*_ = 0.050; BF_exclusion_ = 8.238). Thus, it seems that *Dlg2* haploinsufficiency does not have downstream effects on the expression of related proteins.

### Behaviour in anxiety tests

Figure 3 shows measures from the EPM for time in closed and open arms (3A), head-dips (3B), stretch-attend postures (3C), grooming (3D), distance travelled (3E), velocity (3F) and defecation (3G). Where rodents spent their time in the maze was analysed using repeated measures ANOVA and Bayesian repeated measures ANOVA with within-subjects factor of arm (closed, open) and between-subjects factor of sex and genotype. Ethological measures and movement were analysed with ANOVA and Bayesian ANOVA with factors of sex and genotype. There were no sex effects on any EPM measures as reported in Supplementary Results S.2.1.1.

**Figure 3:**
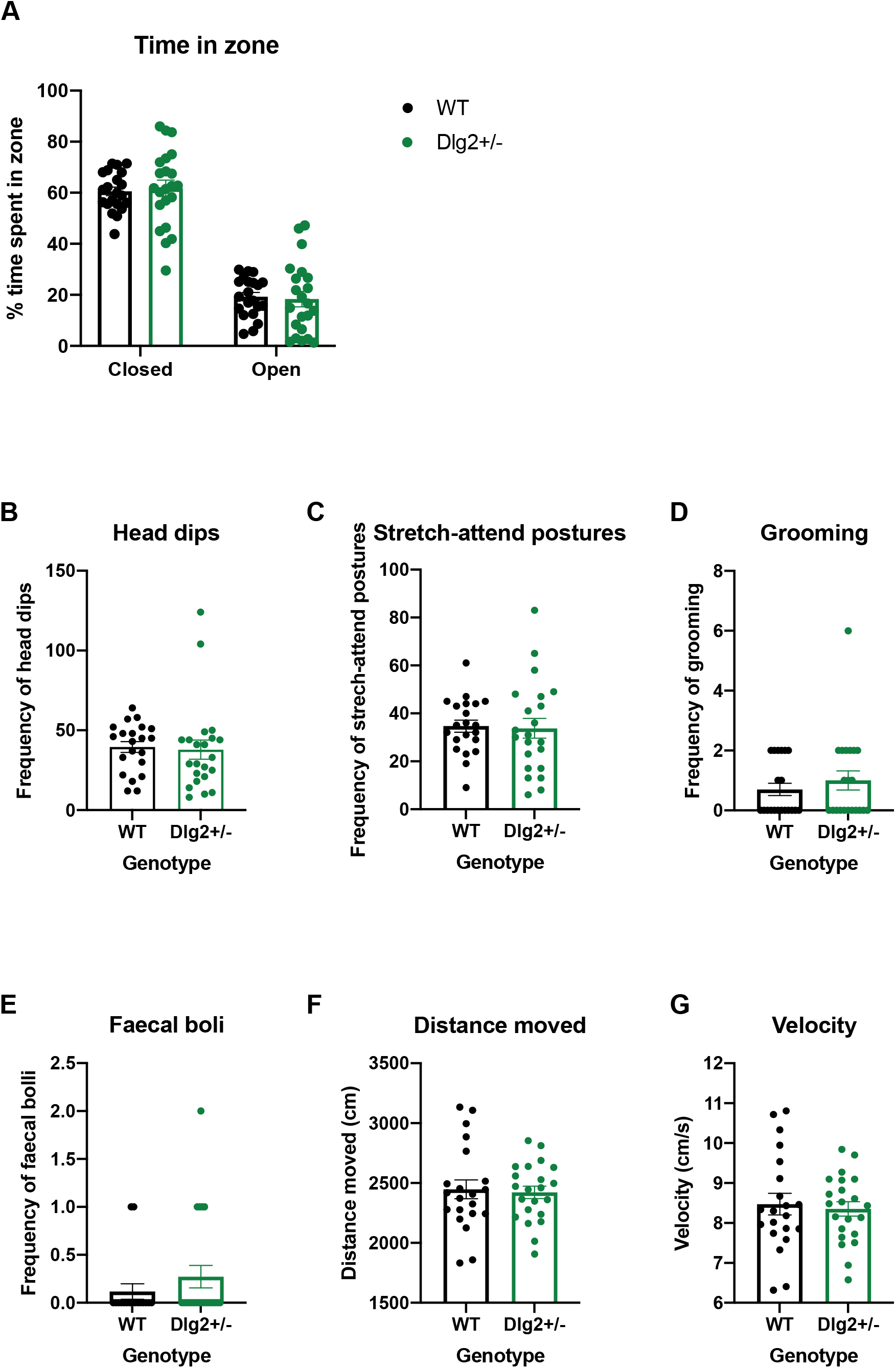
Effect of *Dlg2* heterozygous knockout on anxiety-related behaviour in the elevated plus maze. Data is shown as mean ± SEM with data points representing individuals **A)** time in zone **B)** head dips **C)** stretch-attend postures **D)** grooming **E)** defecation **F)** distance moved **G)** velocity.

All rats showed a tendency to avoid the open arms of the maze however there were no differences between wild-type and *Dlg2*^*+/-*^ rats in the proportion of time spent in open and closed arms: main effect of zone (*F*(1.347, 52.518) = 158.103, *p* < 0.001, *n*^*2*^_*p*_ = 0.802) in the absence of a main effect of genotype (*F*(1, 39) = 1.275, *p* = 0266, *n*^*2*^_*p*_ = 0.032; BF_exclusion_ = 11.197) or zone × genotype interaction (*F*(1.347,52.518) = 0.052, *p* = 0.887, *n*^*2*^_*p*_ = 0.001; BF_exclusion_ = 17.680). There were also no genotype-related difference in the ethological measures assessed: head-dips (*F*(1,33) = 0.034, *p* = 0.854, *n*^*2*^_*p*_ = 0.001; BF_exclusion_ = 4.266), stretch-attend postures (*F*(1,33) = 0.190, *p* = 0.666, *n*^*2*^_*p*_ = 0.006; BF_exclusion_ = 3.528), grooming (*F*(1,33) = 0.002, *p* = 0.961, *n*^*2*^_*p*_ = 0.000; BF_exclusion_ = 3.816) or defecation (*F*(1,39) = 1.505, *p* = 0.227, *n*^*2*^_*p*_ = 0.039; BF_exclusion_ = 2.471). There were also no genotype-related effects on distance travelled (*F*(1,33) = 0.082, *p* = 0.776, *n*^*2*^_*p*_ = 0.002; BF_exclusion_ = 4.182) or velocity (*F*(1,33) = 0.082, *p* = 0.776, *n*^*2*^_*p*_ = 0.002; BF_exclusion_ = 4.151). These findings indicate that *Dlg2*^*+/-*^ rats do not appear to have an anxiety phenotype in the EPM, although both wild-type and *Dlg2*^*+/-*^ rats demonstrated the expected anxiogenic profile for this test.

The distribution of time spent in maze zones was analysed using repeated measures ANOVA and Bayesian repeated measures ANOVA with within-subjects factor of zone (central, outer) and between subjects factor of sex and genotype. Movement and defection were analysed with ANOVA and Bayesian ANOVA with factors of sex and genotype. As with EPM there were no sex effects and these are reported in Supplementary Results S.2.1.2. Figure 4 shows the results from the open field for time in the central and outer areas (4A), velocity (4B), distance moved (4C), and defecation (4D). While there was a general tendency to avoid the central region, there were no differences between wild-type and *Dlg2*^*+/-*^ rats in the proportion of time spent in centre and outer zones: main effect of zone (*F*(1,38) = 9100.383, *p* < 0.001, *n*^*2*^_*p*_ = 0.996) but no main effect of genotype (*F*(1, 38) = 0.697, *p* = 0.409, *n*^*2*^_*p*_ = 0.018; BF_exclusion_ = 7.292) or zone × genotype interaction (*F*(1,38) = 0.232, *p* = 0.633, *n*^*2*^_*p*_ = 0.000; BF_exclusion_ = 6.064). While there was a suggestion that *Dlg2*^*+/-*^ rats defecated more than wild-type controls, there was no significant effect of genotype (*F*(1,38) = 3.769, *p* = 0.060, *n*^*2*^_*p*_ = 0.090; BF_exclusion_ = 0.880). Although the BF was inconclusive here, it should be remembered that there was also no suggestion of a genotype-related effect on defecation in the EPM. There were no genotype differences in either distance travelled (*F*(1,38) = 0.002, *p* = 0.961, *n*^*2*^_*p*_ < 0.001; BF_exclusion_ = 3.551) or velocity (F(1,38) = 0.013, p = 0.908, *n*^*2*^_*p*_ < 0.001; BF_exclusion_ = 3.753). Overall while the experiment demonstrated anxiety generally, with all the rats avoiding the aversive open central region, there were no genotype effects on this nor any other measure in the open field.

**Figure 4:**
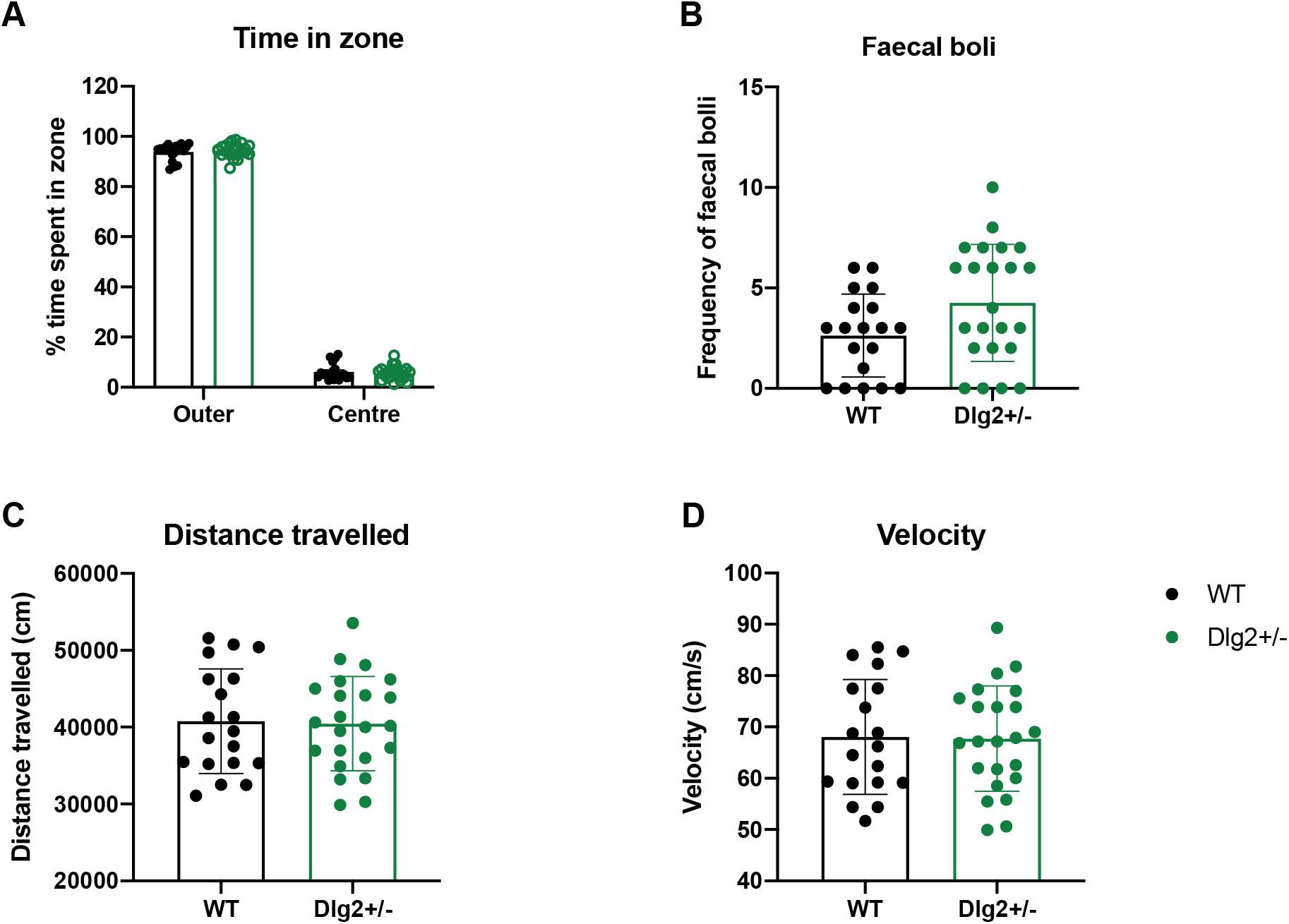
Effect of *Dlg2* heterozygous knockout on open-field measures. Data is shown as mean ± SEM plus individual values for **A)** time in zone **B)** velocity **C)** distance travelled and **D)** defection.

### Acoustic startle response (ASR) and pre-pulse inhibition (PPI)

Response to increasing amplitudes of startle stimuli were assessed using mixed ANOVA and mixed Bayesian ANOVA with within-subjects factor of pulse intensity (70, 80, 90, 100, 110 and 120 dB) and between-subjects factors of genotype and sex from data acquired in the third block of trials from the startle sessions (Figure 5A). There was an absence of sex-related effects on all ASR and PPI measures which are reported in Supplementary Information. There was a significant main effect of pulse intensity (*F*(1.089, 43.542) = 29.705, *p* < 0.001, *n*^*2*^_*p*_ = 0.426) indicating an increased responding at higher startle intensities. There was no significant main effect of genotype (*F*(1, 40) = 0.872, *p* = 0.356, *n*^*2*^_*p*_ = 0.021; BF_exclusion_ = 8.964), nor a genotype × pulse interaction (*F*(1.089, 43.542) = 0.590, *p* = 0.460, *n*^*2*^_*p*_ = 0.015; BF_exclusion_ = 22.407). Thus, wild-type and *Dlg2*^*+/-*^ rats showed equal responding to increasing startle amplitudes, suggesting equivalent acoustic startle responses.

**Figure 5:**
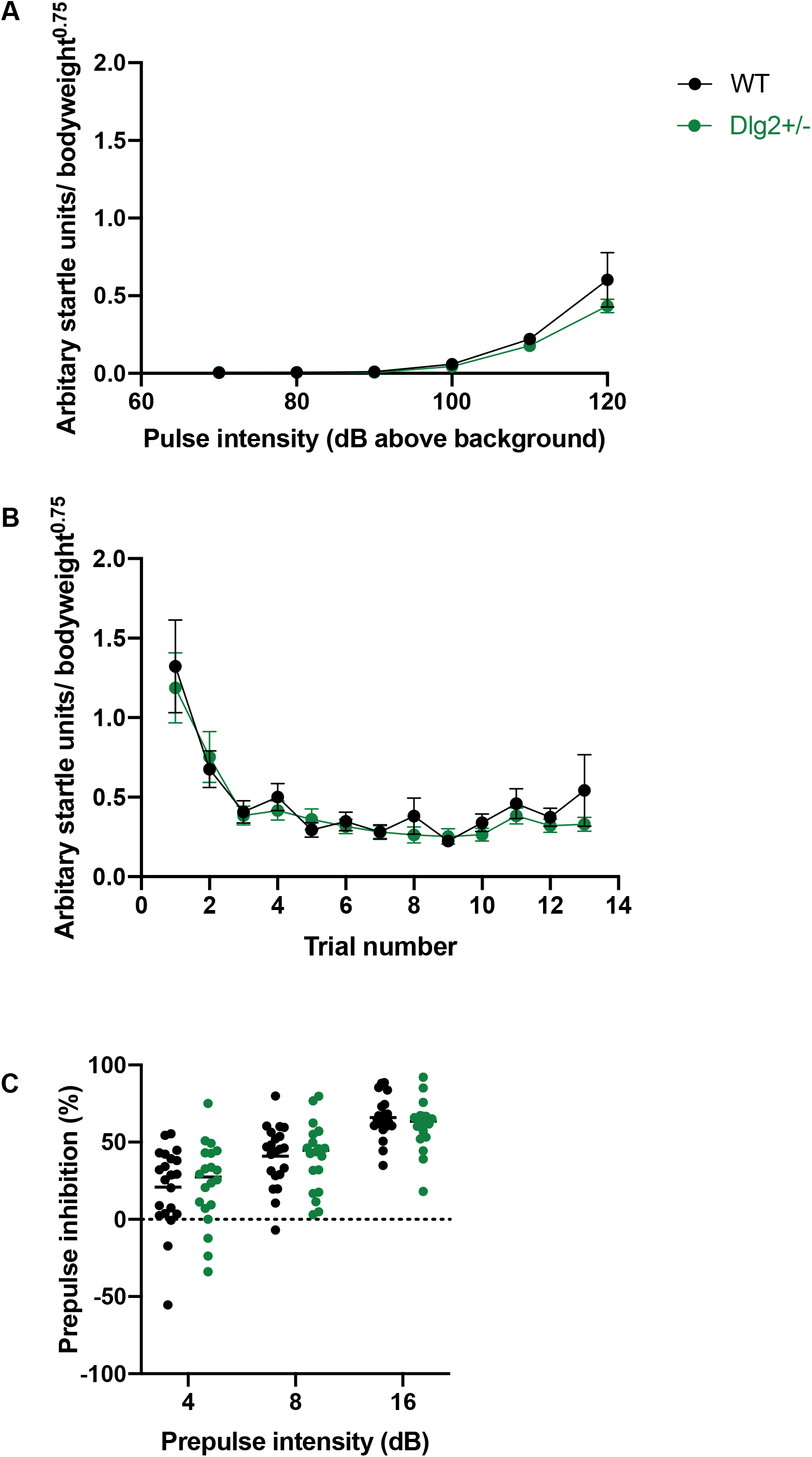
Effect of *Dlg2* heterozygosity on acoustic startle response and pre-pulse inhibition. **A)** Mean ± SEM weight-adjusted ASR to 70-120 dB pulses above background. **B)** Mean ± SEM habituation of startle response through increasing pulse trials and **C)** Mean ± SEM PPI plus individual values with 4, 8 and 16 dB (above background) pre-pulse.

Habituation of the startle response (Figure 5B) to 120db pulse-alone stimuli was assessed using mixed ANOVA and Bayesian mixed ANOVA with within-subjects factors of trial number and between subjects factors of genotype and sex. As expected, the startle response habituated as trials progressed (main effect of trial: *F*(2.130, 85.190) = 13.127, *p* < 0.001, *n*^*2*^_*p*_ = 0.247) but there were no differences between wild-type and *Dlg2*^*+/-*^ rats (genotype × trial interactions: *F*(2.130, 85.190) = 0.498, *p* = 0.621, *n*^*2*^_*p*_ = 0.012; BF_exclusion_ = 643.328 or main effect of genotype: *F*(1, 40) = 0.347, *p* = 0.559, *n*^*2*^_*p*_ = 0.009; BF _exclusion_ = 10.545).

There were also no differences between wild-type and *Dlg2*^*+/-*^ rats on pre-pulse inhibition (Figure 5C). Repeated measures ANOVA and Bayesian repeated measures ANOVA with factors of pre-pulse intensity (4, 8 and 16 dB above background), genotype and sex were used. All rats demonstrated a greater inhibition of the startle response with increasing pre-pulse intensity (significant main effect of pre-pulse: *F*(2, 80) = 83.401, *p* < 0.001, *n*^*2*^_*p*_ = 0.676), but there were no significant genotype × pre-pulse interaction (*F*(2, 80) = 1.097, *p* = 0.339, *n*^*2*^_*p*_ = 0.027; BF_exclusion_ = 5.712 or main effect of genotype (*F*(2, 80) = 0.011, *p* = 0.917, *n*^*2*^_*p*_ = 0.000; BF_exclusion_ = 5.482) on PPI.

### Lick microstructure assessment

Repeated measures ANOVA and Bayesian repeated measures ANOVA were used to analyse consumption and lick cluster data with genotype as a between subject factor and sucrose concentration as a within-subject factor. As Figure 6A shows consumption of sucrose varied with concentration with rats consuming greater volumes of the 16% solution relative to 4% (main effect of concentration, *F*(1,46) = 14.582, *p* = 0.015, *n*^*2*^_*p*_ = 0.121). Genotype had no effect on sucrose consumption as shown by the non-significant main effect of genotype, *F*(1,46) = 0.055, *p* = 0.952, *n*^*2*^_*p*_ = 0.000; BF_exclusion_ = 1.762. The genotype × concentration interaction was significant *F*(1,46) = 7.429, *p* = 0.009, *n*^*2*^_*p*_ = 0.139, reflecting the larger 4-16% based consumption change in the *Dlg2*^*+/-*^ rats compared to the wild-types.

**Figure 6:**
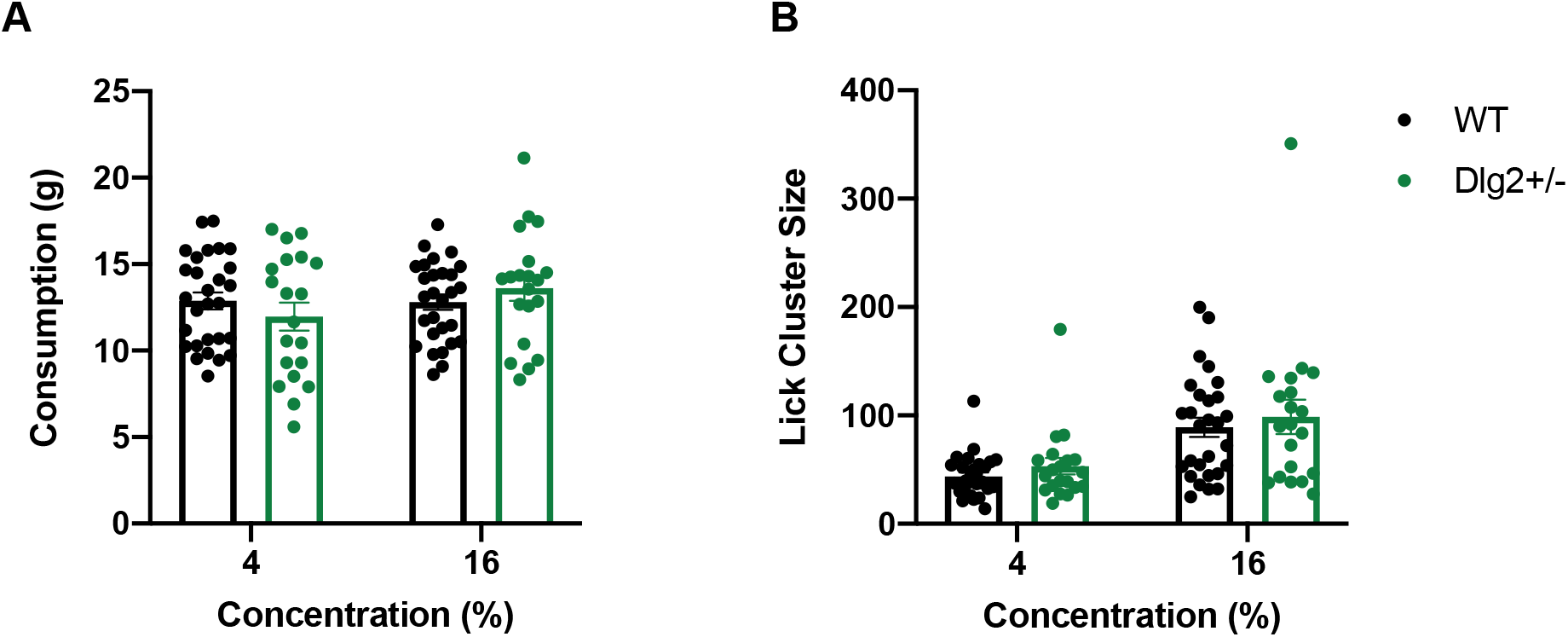
The drinking behaviour of *Dlg2*^*+/-*^ and wild-type rats when presented with low and high concentrations of sucrose. Data is shown as mean ± SEM plus individual values **(A)** consumption (g) and **(B)** lick cluster size.

Figure 6B reveals that lick cluster size also varied with sucrose concentration, as both wild-type and *Dlg2*^*+/-*^ rats showed larger cluster sizes at 16% than to less palatable 4% (main effect of concentration, *F*(1,46) = 56.074, *p* < 0.001, *n*^*2*^_*p*_ = 0.549). Genotype had no effect on lick cluster size at either concentration (non-significant main effect of genotype, *F*(1,46) = 0.642, *p* = 0.427, *n*^*2*^_*p*_ = 0.014 non-significant genotype × concentration interaction *F*(1, 46) = 0.002, *p* = 0.965, *n*^*2*^_*p*_ = 0.000). Evidence for any genotype effect on lick cluster size is inconclusive (BF_exclusion_ = 2.384) as is that for the genotype × concentration interaction (BF_exclusion_ = 2.341). Because evidence for impaired hedonic reactions would require that *Dlg2*^*+/-*^s would have lower lick cluster size than wild-types Bayesian one-tailed independent samples t-tests were done on the lick cluster sizes for 4% and 16% conditions, finding evidence for the absence of this expected *Dlg2*^*+/-*^ less than wild-type effect in both instances (4% BF_01_ = 7.887,16% BF_01_ = 5.735)

The variation in both lick cluster and consumption with concentration is expected and informs that the experiment successfully manipulated the hedonic properties of the stimuli. The lack of any reduction in lick cluster size for the *Dlg2*^*+/-*^ rats suggests there is no suggestion of an anhedonic response to palatable stimuli.

### Social preference test

Raw exploration times for conspecific and object in the social preference test are shown in Figure 7A. These data were analysed by mixed model ANOVA and Bayesian ANOVA with the within-subject factor of item (conspecific, object) and between-subjects factors of sex and genotype. The conspecific was explored more than the object (significant main effect of item:(*F*(1, 50) = 76.012, *p <* 0.001, *n*^*2*^*p* = 0.603) however this did not differ with genotype (non-significant item × genotype interaction, *F*(1,50) = 0.479, *p =* 0.492, *n*^*2*^*p* = 0.009; BF_exclusion_ = 7.678 or genotype main effect *F*(1, 50) = 0.014, *p =* 0.907, *n*^*2*^*p* = 0.000279; BF_exclusion_ = 7.986).

**Figure 7:**
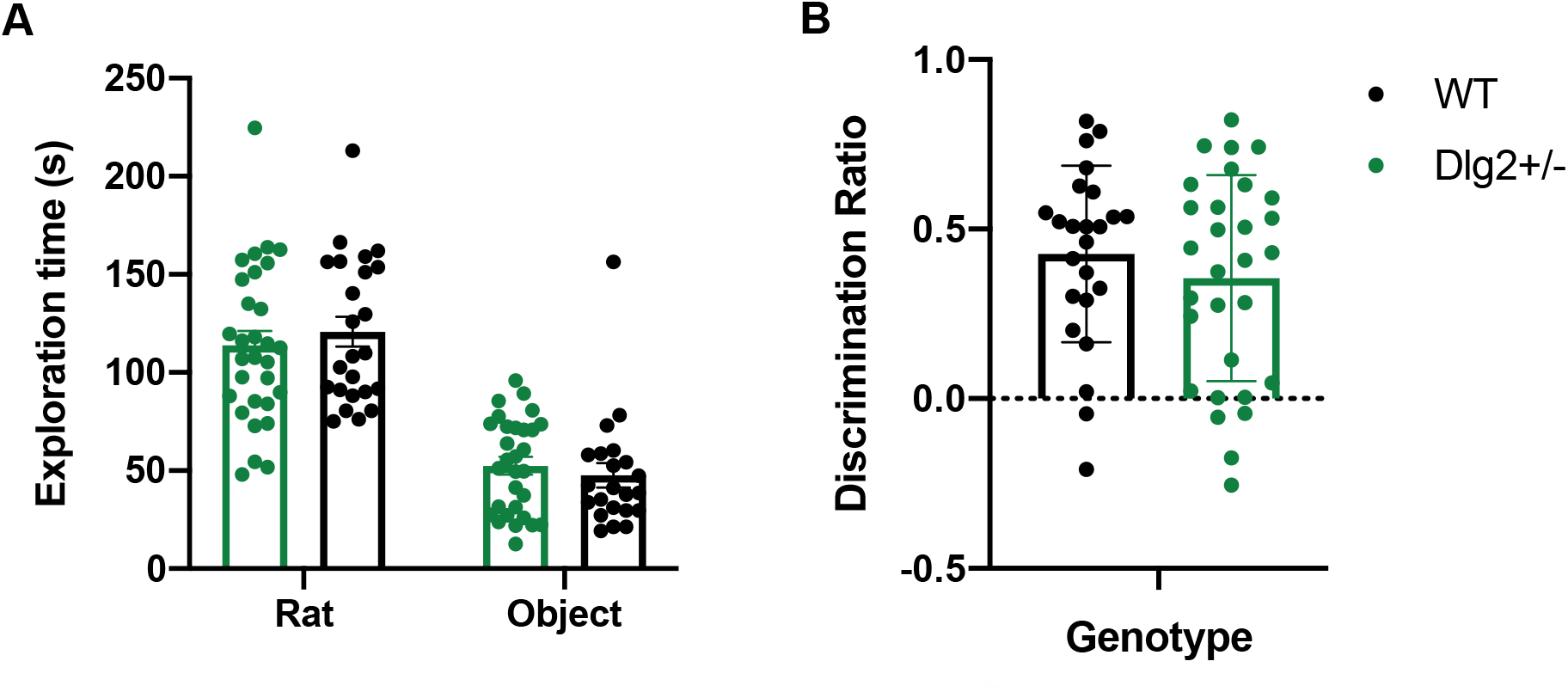
*Dlg2*^*+/-*^ and wild-type exploration times on the social preference task. Data is shown as mean ± SEM plus individual values **A)** raw exploration and **B)** d2 discrimination ratios.

The discrimination ratio for social preference is show in in Figure 7B. This was significantly different from 0 for the entire cohort (one-sample t-test *t(*53*) =* 9.988, *p <* 0.001, *d =* 1.359) reflecting the tendency to explore the conspecific more than the object. There were no genotype differences in discrimination ratio: non-significant main effect of genotype (*F*(1, 50) = 0.850, *p =* 0.361, *n*^*2*^*p* = 0.017; BF_exclusion_ = 4.973). This reflects the fact that rats explored the conspecific more than the object as expected, yet *Dlg2* haploinsufficiency has no influence on this tendency.

### PCP-induced locomotion

Figure 8 shows the distance travelled in the arena over the 120-minute test period (30 mins preceding 5mg/kg PCP administration and the following 90 mins) for wild-type and *Dlg2*^*+/-*^ rats. Distance travelled was analysed separately for the 30 minutes preceding injection and the 90 minutes post injection using repeated measures ANOVA with the repeated measures factor of time bin (3 × 10-minute bins covering the 30 minutes preceding injection and 9 × 10-minute bins 90 minutes post-injection) and between-subjects factors of sex and genotype.

**Figure 8:**
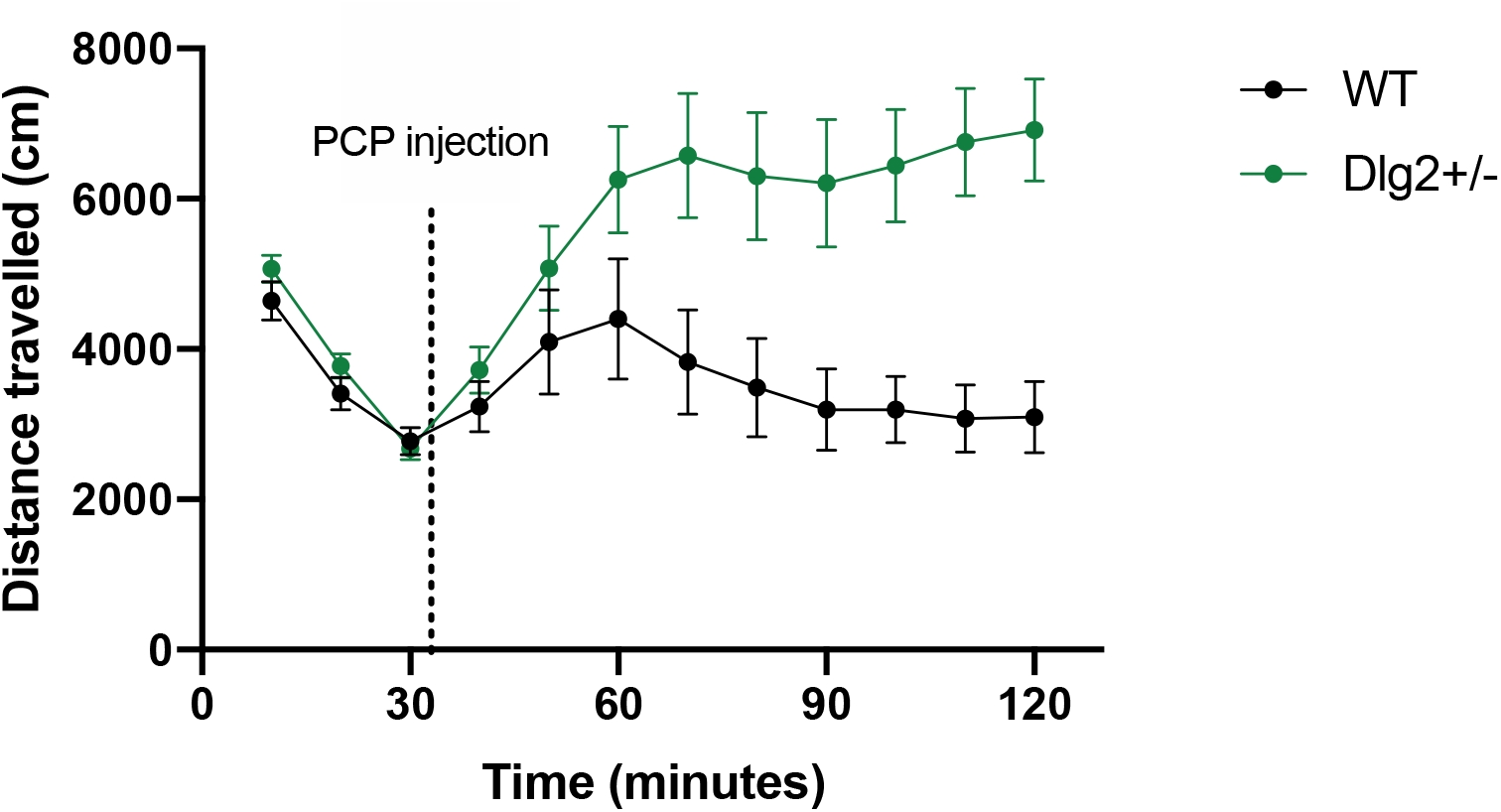
Locomotor activity in response to PCP injection in *Dlg2*^*+/-*^ and wild-type rats. Data is shown as mean ± SEM distance travelled plotted in 10-minute bins. The dotted line just past 30 minutes denotes when the PCP injection occurred.

When analysing distance travelled for the 30 minutes before the injection of 5mg/kg PCP, there was a main effect of time bin (*F*(2, 98) = 227.098, *p <* 0.001, *n*^*2*^_*p*_ = 0.823) as activity decreased through rats initial habituation. There was no main effect of genotype (*F*(1, 49) = 0.452, *p* = 0.505, *n*^*2*^_*p*_ = 0.009; BF_exclusion_ = 2.165) but the time bin × genotype interaction approached significance (*F*(2, 98) = 2.516, *p* = 0.086, *n*^*2*^_*p*_ = 0.049; BF_exclusion_ = 1.243) with *Dlg2*^*+/-*^ rats showing a slight tendency to a faster reduction in activity across the 30 minute pre-injection habituation period. At 30 minutes there was no difference in activity between genotypes (*t*(51) = −0.420, *p* = 0.676, *d* = −0.116; BF_01_ = 3.171), providing an equivalent baseline before PCP injection. Comparisons for genotype at 10 minutes (*t*(52) = 1.375, *p* = 0.175, *d* = 0.375; BF_01_ = 2.359) and 20 minutes (*t*(51) = 1.375, *p* = 0.175, *d* = 0.379; BF_01_ = 1.721) were also non-significant.

Following injection of PCP, there was a significant increase in movement by both wild-type and *Dlg2*^*+/-*^ rats (main effect of time bin: *F*(2.829, 141.460) = 4.044, *p* = 0.010, *n*^*2*^_*p*_ = 0.075), although the effect was much greater and longer lasting in *Dlg2*^*+/-*^ rats, as demonstrated by a significant time bin × genotype interaction (*F*(2.829, 141.460) = 5.125, *p* = 0.003, *n*^*2*^_*p*_ = 0.093) and main effect of genotype (*F*(1, 50) = 9.873, *p* = 0.003, *n*^*2*^_*p*_ = 0.165). Follow up analyses of the time bin by genotype interaction indicated that the *Dlg2*^*+/-*^ animals were more active than wild-types on time bin 70 and onwards (smallest *t*(56) = 2.466, *p* = 0.017, *d* = 0.651) but not on time bin 60 and before (largest *t*(56) = 1.738, *p* = 0.088, *d* = 0459).

## Discussion

This work presents the characterisation of which molecular and behavioural capabilities are spared and impaired in the *Dlg2* heterozygous rat model; a model with direct clinical relevance to CNVs that increase risk for a variety of psychiatric conditions including schizophrenia ^5^, autism ^3^ and intellectual disability ^4^. The model is specific to *Dlg2* and valid, with evidence that mRNA and protein expression of *Dlg2* is reduced in the absence of changes to levels of other *Dlgs*. Behaviourally *Dlg2*^*+/-*^ rats performed comparably to wild-types on tests of anxiety, hedonic reactions, social behaviour, and sensorimotor gating.

When locomotor response to PCP challenge was assessed, *Dlg2*^*+/-*^ rats demonstrated a potentiated response to the drug. This demonstrates the first behavioural correlate of *Dlg2* heterozygosity and is in line with electrophysiological data demonstrating a change in NMDAR function in *Dlg2* heterozygous rats ^29^. This contrasts with work on *Dlg2* homozygous knockdown models where no change in NMDAR function has been documented ^43–45^, however Zhang et al ^22^ observed that PSD-93 deficiency in cortical neurons reduces the expression of NR2A and NR2B NMDAR subunits and changes Ca^2+^ influx through NMDARs.

Altered psychostimulant sensitivity has also been shown in other psychosis-relevant CNV rodent models, the 22q11.2 ^46^ and 1q21.1 ^47^ microdeletion mouse models. In the 22q11.2 mouse this manifested as an exaggerated locomotor response to PCP and ketamine. In the 1q21.1 mouse exaggerated locomotor behaviour was seen in response to amphetamine but was not significant with administration of PCP, yet PCP resulted in sensorimotor gating impairments. As administration of non-competitive NMDAR antagonists in healthy rodents and humans can induce a behavioural syndrome isomorphic to positive and negative symptoms of schizophrenia ^48–50^ and exacerbates positive and negative symptoms in schizophrenic patients ^51–53^ the finding of PCP sensitivity in these rodent CNV models may highlight a general psychosis susceptibility. Findings such as these contribute to the hypoglutamaterigic hypothesis of psychosis ^54^. However, it is difficult to use these findings to determine which biological processes are altered in CNV rodent models. Acute PCP administration has been reported to activate serotonergic, glutamatergic, noraderenergic, cholinergic and neurotensinergic transmission in rodents and monkeys ^55–57^. It will be informative to focus on investigating how individual behavioural alterations to NMDAR antagonists, and their timescales, may be subserved by particular neurotransmitter dysfunctions to identify mechanisms underpinning these in genetic disorder models.

Coherence between findings on homozygous mouse models, heterozygous mouse models and the CRISPR-Cas9 generated *Dlg2*^*+/-*^ rat were mixed. Findings of impaired social preference, increased grooming and hypoactivity in the open field test of anxiety in homozygous mouse models ^1^ did not replicate in the *Dlg2*^*+/-*^ rat. Comparisons between complete and heterozygous gene knockout models allow a distinction to be made between knowledge about the function of a protein and processes which require complete PSD93 levels. The difference here implies that having some functional PSD93 might ‘rescue’ these phenotypes. This could be supported by PSD93 acting in tandem with PSD95 or other MAGUKs. Where social behaviour is concerned it has been shown that there is a similarity in social deficits in *PSD95*^*+/-*^ mice and *PSD93*^*-/-*^ mice, with increased expression of PSD93 in the hippocampus of *PSD95*^*+/-*^ mice implying that PSD93 is acting to compensate in this mouse model ^17^. In the *Dlg2*^*+/-*^ rat it could be that intact PSD95 in the presence of some PSD93 was sufficient to support intact social preference performance, meaning that while PSD93 has some role in social behaviours it is not so essential that genetic haploinsufficiency produces a gross deficit.

The observed lack of anxiety and pre-pulse inhibition phenotypes in the *Dlg2*^*+/-*^ rat were also found in *Dlg2*^*+/-*^ mice^58^. However, *Dlg2*^*+/-*^ mice demonstrated a subtle deficit of habituation to the acoustic stimulus in this facet of the sensorimotor gating task, which was not seen in the *Dlg2*^*+/-*^ rat. This may be due to differences in model generation, with the heterozygous mouse generated by introduction of a cassette upstream of the critical exon (14) on chromosome 7 while the heterozygous rat was generated by 7bp deletion within the rat *Dlg2* gene resulting in a frame shift and premature stop codon. This behavioural difference also points to potential caveats in comparisons across psychiatric risk models with different species backgrounds, which is also relevant when comparing homozygous mice with the rat model.

Another point of interest is the lack of interactions between sex and genotype in this work (see Supplement 1 for detailed data), which in the main was conducted on mixed sex cohorts. This is an important inclusion in the rodent modelling of psychiatric susceptibility literature which is often conducted on male-only cohorts, and thus runs the risk of limited translational generalisability. Critically, the results here relating to the genotype manipulation were unaffected by the sex of the animals.

## Conclusion

The *Dlg2*^*+/-*^ rat validly models a single copy deletion of *Dlg2* including concomitant mRNA and protein decrease in the absence of obvious compensation. No gross behavioural deficits on tasks relevant to a broad spectrum of psychiatric phenotypes were found, except for exaggerated hyperlocomotion in response to PCP, an NMDAR-antagonist. A similar selectivity in phenotype is seen for other psychiatric CNV models such as the 1q21.1 mouse ^47^. This demonstrates the behavioural subtlety of the model and highlights issues with drawing clinical conclusions from homozygous models. It also paves the way for investigation into more complex behavioural domains such as memory and learning without the concern of confounds from anxiety, hedonic processing, hearing, and social processing.

## Acknowledgements

Research funded by the Wellcome Trust Strategic Award ‘DEFINE’ (100202/Z/12/Z) to RP and the Wellcome Trust studentship for four-year PhD programmes in Science (BV17108004) to SW.

