## Supplementary Information for "Behavioural and molecular characterisation of the Dlg2 haploinsufficiency rat model of genetic risk for psychiatric disorder"

Data Supplement

**Running title**

SUPPLEMENTAL DATA – SEX EFFECTS IN THE CHARACTERISATION OF THE DLG2 +/- RAT

### Supplementary Results

#### S1 Example Western blot

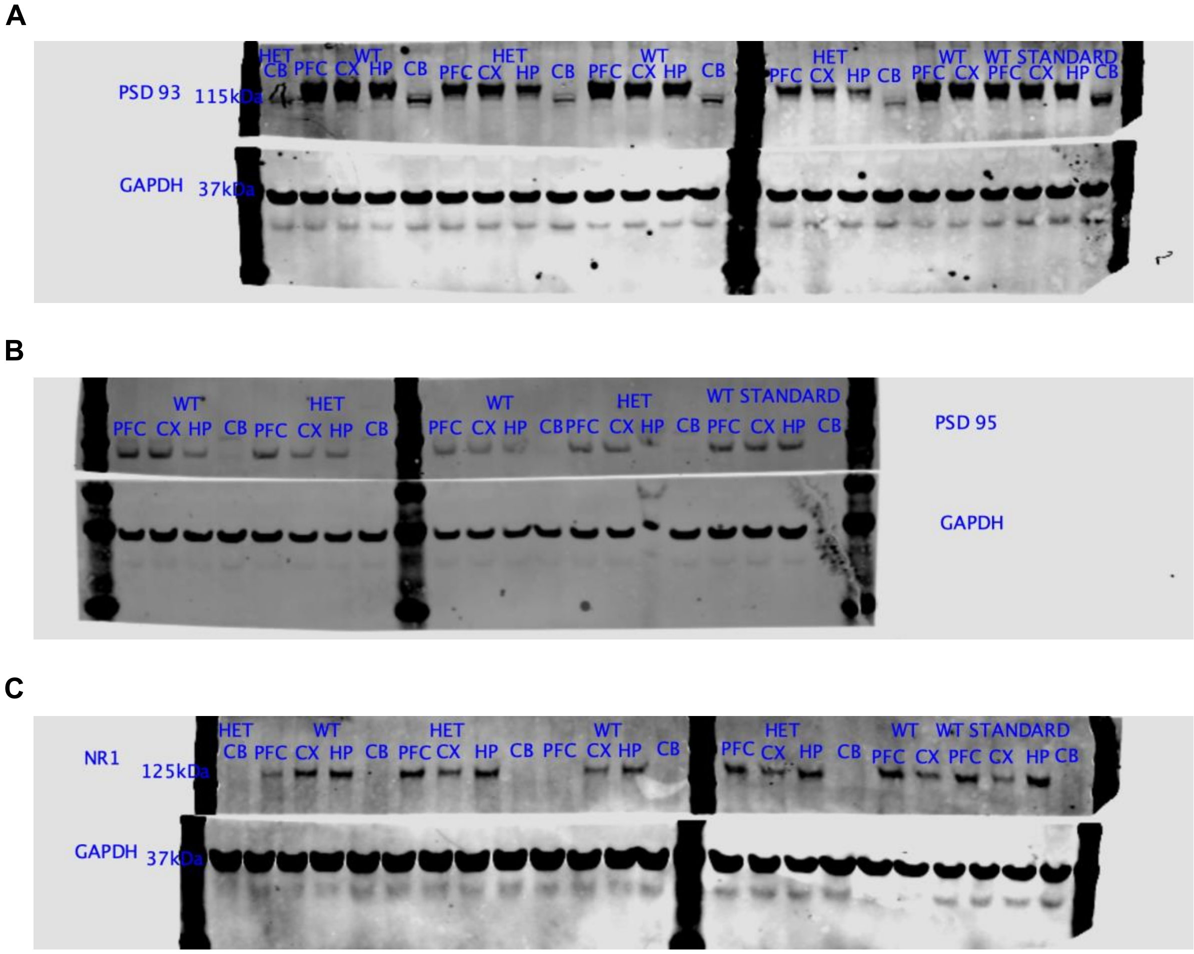


**Figure S1: Sample blots showing the same samples and brain regions for PSD93 (A), PSD95 (B) and NR1 (C).** Lack of bands for NR1 cerebellum shown.

#### S2 Sex effects in the characterisation of the *Dlg2*$^{+/-}$ rat

The use of male-only cohorts when investigating psychiatric risk with rodent models is prevalent in the literature but this may result in conclusions that overlook sex differences in how these risk factors operate. Thus, the current experimental work used both male and female animals to allow sex differences to be investigated – however, as noted in the main text, no sex by genotype interactions were observed for any of the outcome measures considered. Thus, the main text report focused on genotype effects regardless of sex, while here we report the full results of the analyses including sex as a variable (oestrus stage was included as a variable initially – but had no effects so was not reported here). These support the conclusion that the current results are consistent with the *Dlg2*$^{+/-}$ manipulation being similar (or similarly absent) in male and female rats.

### S2.1 Behaviour on anxiety tests

#### S2.1.1 Elevated Plus Maze

As shown in Figure S1 sex had no effect on any measures recorded in the EPM including time in closed and open arms (Figure S2A), head dips (Figure S2B), stretch-attend postures (Figure S2C), grooming (Figure S2D), distance travelled (Figure S2E), velocity (Figure S2G) or defecation (Figure S2F). All main effects and interactions (along with Bayes factors providing evidence for the null effects) in the EPM assays can be seen in Table S1.

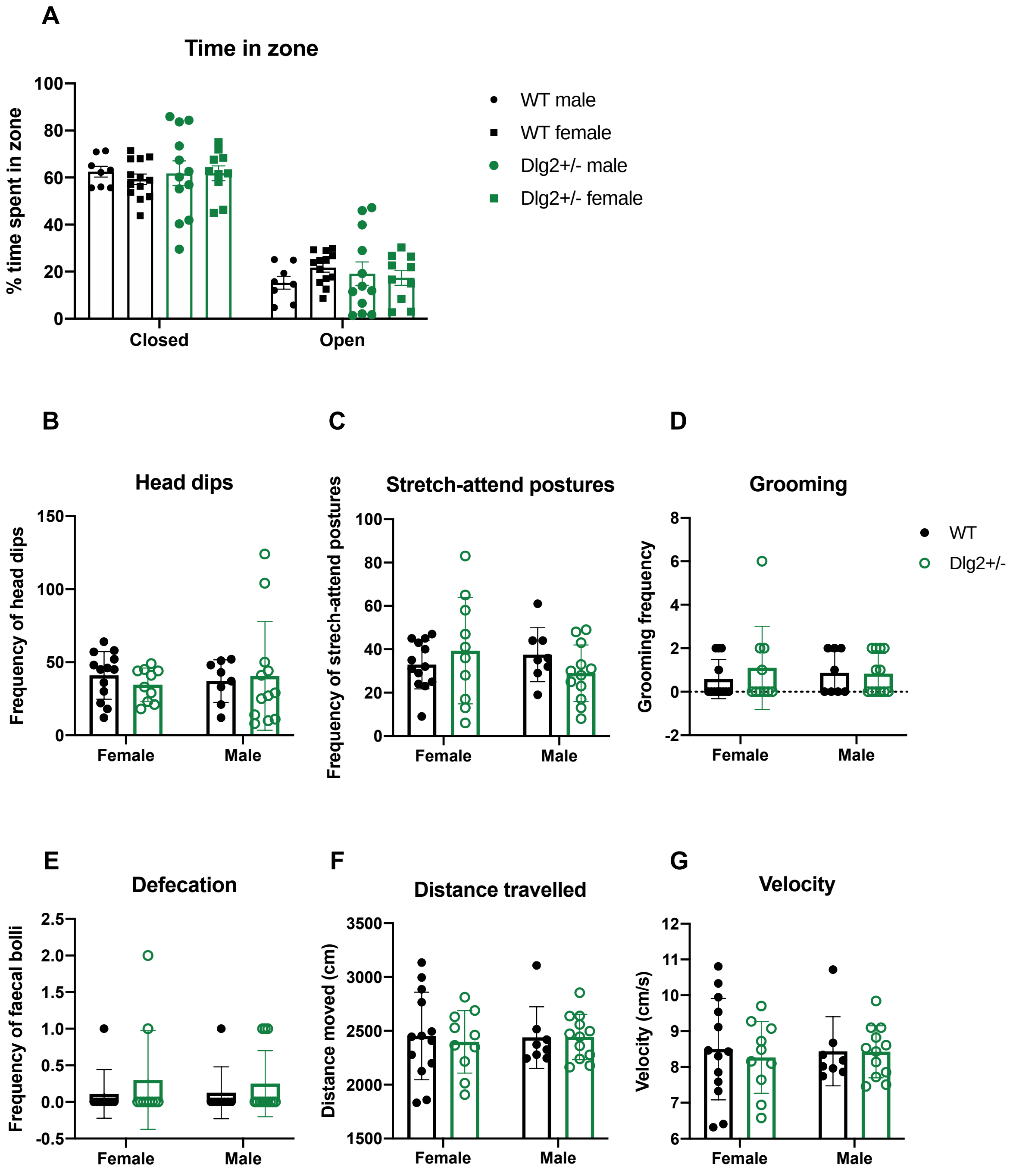

**Figure S2: Effect of *Dlg2* heterozygous knockout and sex on anxiety-related behaviour in the EPM.** Mean ± SEM with data points representing individuals **A)** time in zone **B)** head dips **C)** stretch-attend postures **D)** grooming **E)** defecation **F)** distance moved **G)** velocity.

Table S1: Repeated measures ANOVA and Bayesian ANOVA inferential statistics for analyses including sex as a factor on EPM measures.

| Analysis | Effect | *F* | *p* | $n^2_p$ | $BF_{exclusion}$ |
|---|---|---|---|---|---|
| Time in zone | Zone main effect | (1.347, 52.518) = 158.103 | < 0.001 | 0.802 | 0.000 |
| | Genotype main effect | (1,39) = 1.275 | 0.266 | 0.032 | 11.197 |
| | Sex main effect | (1,39) = 1.497 | 0.228 | 0.037 | 10.610 |
| | Genotype × sex | (1,39) = 0.261 | 0.612 | 0.007 | 40.913 |
| | Genotype × zone | (1.347,52.518) = 0.052 | 0.887 | 0.001 | 17.680 |
| | Zone × sex | (1.347,52.518) = 0.277 | 0.670 | 0.007 | 14.764 |
| | Zone × genotype × sex | (1.347,52.518) = 0.843 | 0.395 | 0.021 | 539.677 |
| Head dips | Genotype main effect | (1,33) = 0.034 | 0.854 | 0.001 | 4.266 |
| | Sex main effect | (1,33) = 0.054 | 0.818 | 0.002 | 4.222 |
| | Genotype × sex | (1,33) = 0.107 | 0.746 | 0.003 | 9.677 |
| Stretch attend postures | Genotype main effect | (1,33) = 0.190 | 0.666 | 0.006 | 3.528 |
| | Sex main effect | (1,33) = 0.088 | 0.769 | 0.003 | 3.578 |
| | Genotype × sex | (1,33) = 1.493 | 0.230 | 0.043 | 5.343 |
| Grooming | Genotype main effect | (1,33) = 0.002 | 0.961 | 0.000 | 3.816 |
| | Sex main effect | (1,33) = 0.001 | 0.970 | 0.000 | 3.825 |
| | Genotype × sex | (1,33) = 0.748 | 0.393 | 0.022 | 7.897 |
| Distance travelled | Genotype main effect | (1,33) = 0.082 | 0.776 | 0.002 | 4.182 |
| | Sex main effect | (1,33) = 0.092 | 0.764 | 0.003 | 4.170 |

| | | | | | |
|---|---|---|---|---|---|
| | Genotype × sex | (1,33) = 0.003 | 0.995 | 0.000 | 10.242 |
| Velocity | Genotype main effect | (1,33) = 0.082 | 0.776 | 0.002 | 4.151 |
| | Sex main effect | (1,33) = 0.092 | 0.763 | 0.003 | 4.139 |
| | Genotype × sex | (1,33) = 0.003 | 0.995 | 0.000 | 9.743 |
| Defecation | Genotype main effect | (1,39) = 1.505 | 0.227 | 0.039 | 2.471 |
| | Sex main effect | (1,39) = 0.00000460 | 0.995 | 0.002 | 3.970 |
| | Genotype × sex | (1,39) = 0.120 | 0.731 | 0.008 | 5.940 |

#### S2.1.2 Open Field

Sex also had no effect on open-field hyperactivity and anxiety measures as shown in Figure S2, including time in centre and peripheral zones (Figure S3A), velocity (Figure S3B), distance travelled (Figure S3C) and defecation (Figure S3D). Inferential statistics including sex for these measures can be found in Table S2, which shows significant sex effects for distance travelled and velocity in the open field yet no sex × genotype interactions.

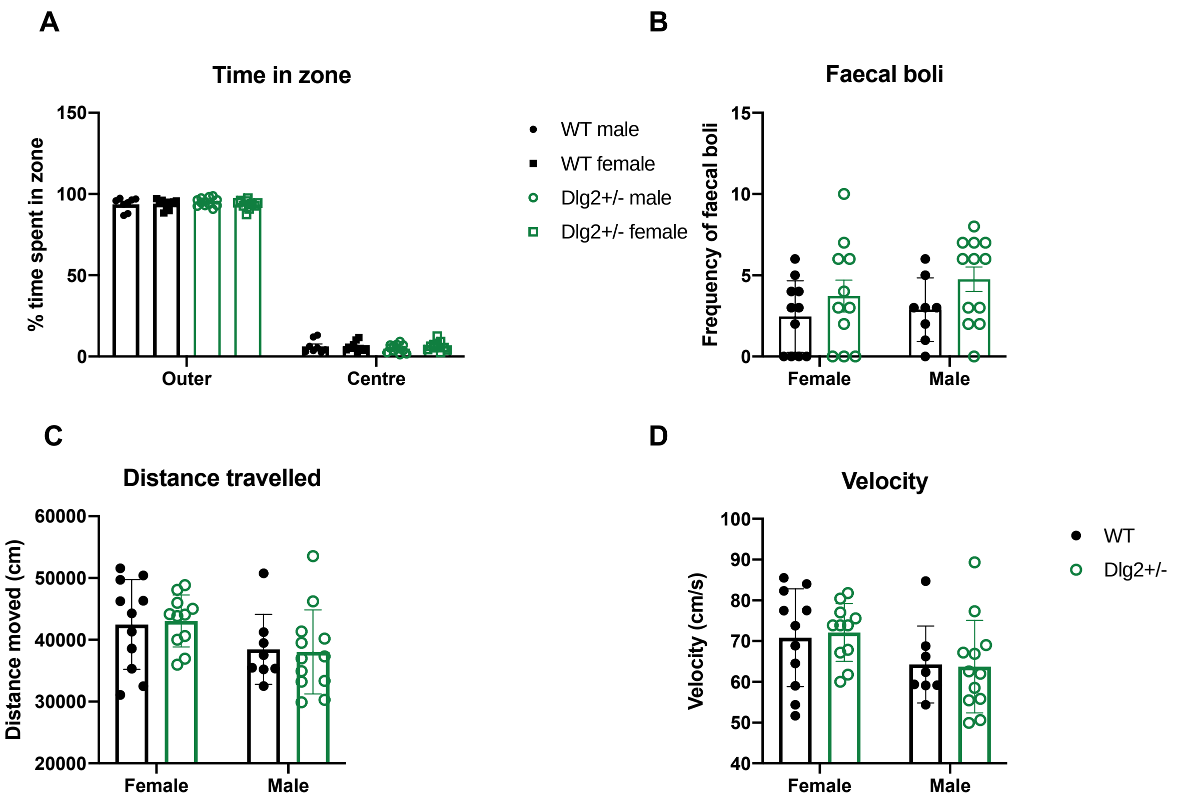

**Figure S3: Effect of *Dlg2* heterozygous knockout and sex on open-field measures A)** time in zone **B)** defection **C)** distance travelled and **D)** velocity. Mean ± SEM with data points representing individuals.

Table S2: Repeated measures ANOVA and Bayesian ANOVA inferential statistics for sex effects and interactions on open field measures.

| Analysis | Effect | *F* | *p* | $n^2_p$ | $BF_{exclusion}$ |
|---|---|---|---|---|---|
| Time in zone | Zone main effect | (1,38) = 9100.383 | < 0.001 | 0.996 | 0.000 |
| | Genotype main effect | (1,38) = 0.697 | 0.409 | 0.018 | 7.292 |
| | Sex main effect | (1,38) = 0.222 | 0.641 | 0.006 | 7.169 |
| | Genotype × sex | (1,38) = 0.095 | 0.759 | 0.003 | 19.332 |
| | Zone × genotype | (1,38) = 0.232 | 0.633 | 0.000 | 6.064 |
| | Zone × sex | (1,38) = 0.362 | 0.551 | 0.000 | 4.372 |
| | Zone × genotype × sex | (1,38) = 0.973 | 0.330 | 0.000 | 42.124 |
| Velocity | Genotype main effect | (1,38) = 0.013 | 0.908 | 0.000 | 3.753 |
| | Sex main effect | (1,38) = 5.481 | 0.025 | 0.126 | 0.453 |
| | Genotype × sex | (1,38) = 0.083 | 0.775 | 0.002 | 3.462 |
| Distance travelled | Genotype main effect | (1,38) = 0.002 | 0.961 | 0.000 | 3.551 |
| | Sex main effect | (1,38) = 5.526 | 0.024 | 0.127 | 0.434 |
| | Genotype × sex | (1,38) = 0.061 | 0.806 | 0.002 | 3.853 |
| Defecation | Genotype main effect | (1,38) = 3.769 | 0.060 | 0.090 | 0.880 |
| | Sex main effect | (1,38) = 0.792 | 0.379 | 0.020 | 2.601 |
| | Genotype × sex | (1,38) = 0.138 | 0.712 | 0.004 | 3.300 |

#### S2.2 Sensorimotor gating

Rats of different sexes performed comparably on tests of startle to increasing auditory stimuli (Figure S4A), habituation of startle response at 105 dB (Figure S4B) and 120 dB (Figure S4C) and pre-pulse inhibition at 105 dB (Figure S4D) and 120 dB (Figure S4E). All main effects and interactions (along with Bayes factors providing evidence for the null effects) in the EPM assays can be seen in Table S3.

A

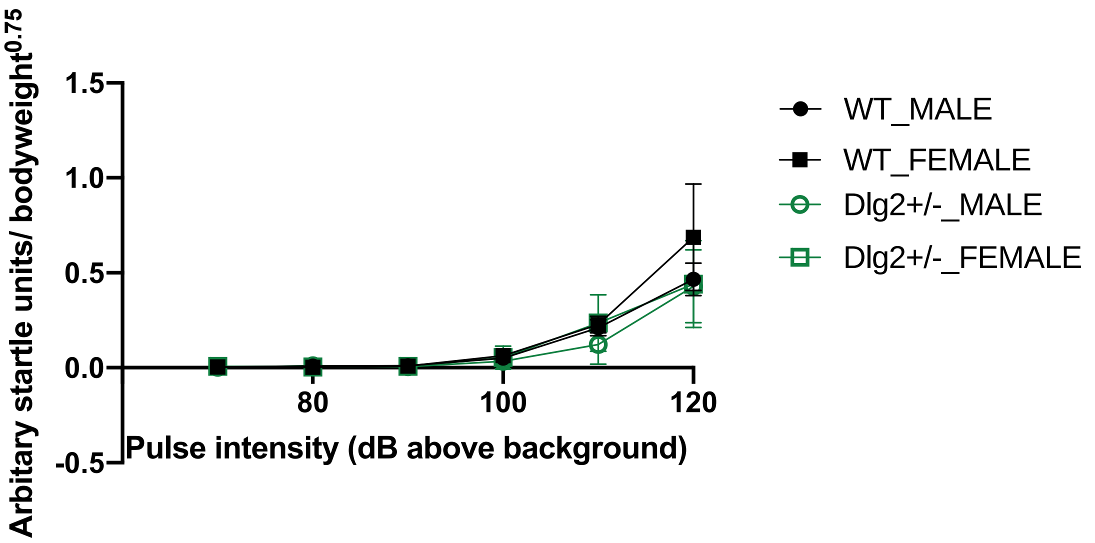
Arbitary startle units/ bodyweight$^{0.75}$
1.5
1.0
0.5
0.0
-0.5
80
100
120
Pulse intensity (dB above background)
WT_MALE
WT_FEMALE
Dlg2+/-_MALE
Dlg2+/-_FEMALE

B

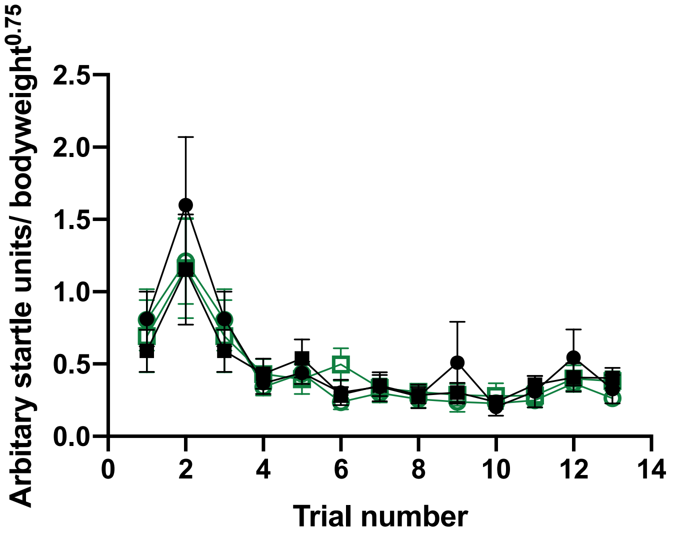
Arbitary startle units/ bodyweight$^{0.75}$
2.5
2.0
1.5
1.0
0.5
0.0
0
2
4
6
8
10
12
14
Trial number

C

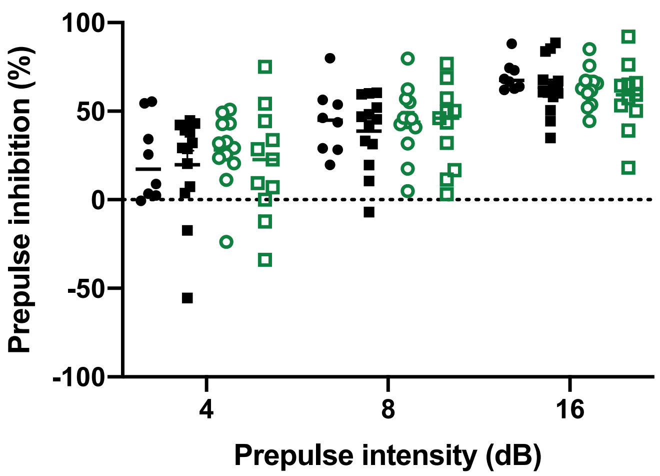
Prepulse inhibition (%)
100
50
0
-50
-100
4
8
16
Prepulse intensity (dB)

**Figure S4: Effect of *Dlg2* heterozygosity and sex on acoustic startle response and pre-pulse inhibition. A)** Mean ± SEM weight-adjusted ASR to 70-120 dB pulses above background. **B)** Habituation of startle response through increasing pulse trials. Mean ± SEM ASR shown at 120 dB and **C)** 105 dB. **D)** Mean ± SEM with data points representing individuals PPI by a 4, 8 and 16 dB (above background) pre-pulse on 120 dB pulse and **E)** 105 dB pulse.

Table S3: Repeated measures ANOVA and Bayesian ANOVA inferential statistics for sex effects and interactions on sensorimotor gating measures.

| Analysis | Effect | *F* | *p* | $n^2_p$ | $BF_{exclusion}$ |
|---|---|---|---|---|---|
| Weight-adjusted ASR to 70-120 dB pulses above background | Pulse main effect | (1.089, 43.542) = 29.705 | < 0.001 | 0.426 | < 0.0001 |
| | Genotype main effect | (1, 40) = 0.872 | 0.356 | 0.021 | 8.964 |
| | Sex main effect | (1, 40) = 0.858 | 0.360 | 0.021 | 9.573 |
| | Genotype × sex | (1, 40) = 0.056 | 0.815 | 0.001 | 38.490 |
| | Genotype × pulse | (1.089, 43.542) = 0.590 | 0.460 | 0.015 | 22.407 |
| | Sex × pulse | (1.089, 43.542) = 0.450 | 0.522 | 0.011 | 37.906 |
| | Genotype × sex × pulse | (1.089, 43.542) = 0.485 | 0.505 | 0.012 | 9759.370 |
| Habituation of startle response (105 dB) | Trial main effect | (1.573, 62.934) = 6.058 | 0.007 | 0.132 | < 0.0001 |
| | Sex main effect | (1, 40) = 2.127 | 0.153 | 0.050 | 4.607 |
| | Genotype main effect | (1, 40) = 1.497 | 0.228 | 0.036 | 6.416 |
| | Genotype × sex | (1, 40) = 0.056 | 0.813 | 0.001 | 18.698 |
| | Trial × genotype | (1.573, 62.934) = 0.519 | 0.555 | 0.013 | 127.064 |
| | Trial × sex | (1.573, 62.934) = 0.681 | 0.476 | 0.017 | 102.095 |

| | | | | | |
|---|---|---|---|---|---|
| | Trial × sex × genotype | (1.573, 62.934) = 1.247 | 0.288 | 0.030 | 63776.307 |
| Habituation of startle response (120 dB) | Trial main effect | (2.130, 85.190) = 13.127 | < 0.001 | 0.247 | < 0.0001 |
| | Sex main effect | (1, 40) = 0.012 | 0.913 | 0.000 | 10.694 |
| | Genotype main effect | (1, 40) = 0.347 | 0.559 | 0.009 | 10.545 |
| | Genotype × sex | (1, 40) = 0.072 | 0.790 | 0.002 | 32.699 |
| | Trial × genotype | (2.130, 85.190) = 0.498 | 0.621 | 0.012 | 643.328 |
| | Trial × sex | (2.130, 85.190) = 1.160 | 0.320 | 0.028 | 174.990 |
| | Trial × sex × genotype | (2.130, 85.190) = 0.543 | 0.594 | 0.013 | 5964000 |
| PPI 105 dB | Pre-pulse main effect | (1.380, 55.210) = 42.203 | < 0.001 | 0.513 | < 0.0001 |
| | Sex main effect | (1, 40) = 1.495 | 0.229 | 0.036 | 6.523 |
| | Genotype main effect | (1, 40) = 0.696 | 0.409 | 0.017 | 3.437 |
| | Genotype × sex | (1, 40) = 1.106 | 0.299 | 0.027 | 8.717 |
| | Pre-pulse × genotype | (1.380, 55.210) = 0.314 | 0.650 | 0.008 | 3.713 |
| | Pre-pulse × sex | (1.380, 55.210) = 0.104 | 0.827 | 0.003 | 7.207 |
| | Pre-pulse × genotype × sex | (1.380, 55.210) = 1.952 | 0.163 | 0.047 | 87.336 |
| PPI 120 dB | Pre-pulse main effect | (2, 80) = 83.401 | < 0.001 | 0.676 | < 0.0001 |
| | Sex main effect | (1, 40) = 0.902 | 0.348 | 0.022 | 4.447 |
| | Genotype main effect | (1, 40) = 0.011 | 0.917 | 0.000 | 5.482 |

| | | | | | |
|---|---|---|---|---|---|
| | Genotype × sex | (1, 40) = 0.0001929 | 0.989 | 0.000 | 9.240 |
| | Pre-pulse × sex | (2, 80) = 0.030 | 0.970 | 0.001 | 10.064 |
| | Pre-pulse × genotype | (2, 80) = 1.097 | 0.339 | 0.027 | 5.712 |
| | Pre-pulse × genotype × sex | (2, 80) = 0.177 | 0.838 | 0.004 | 206.171 |

#### S2.3 Social preference

There were no sex effects on exploration in the social preference task as seen for raw exploration time (Figure S5A) and discrimination ratio (Figure S5B). All main effects and interactions (along with Bayes factors providing evidence for the null effects) in the EPM assays can be seen in Table S4.

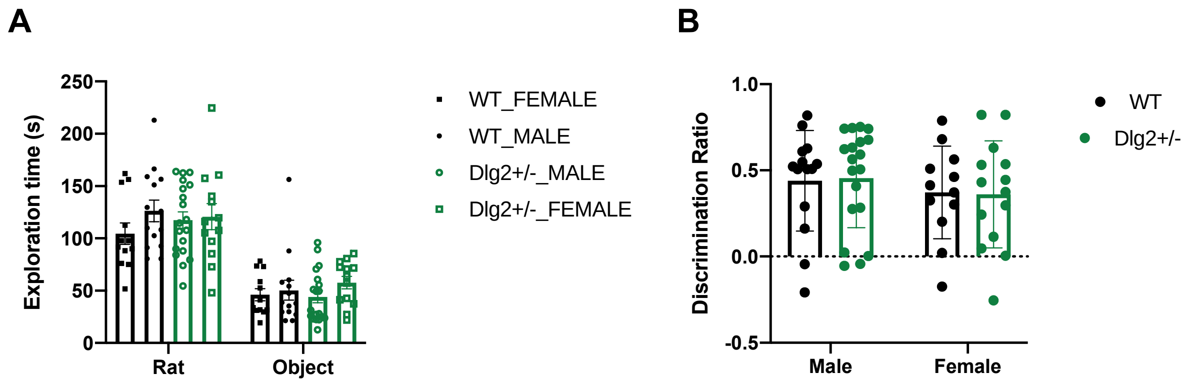


**Figure S5: Effect of *Dlg2* heterozygosity and sex on the social preference task.** Mean and SEM ± with data points representing individuals **A)** raw exploration and **B)** d2 discrimination ratios.

Table S4: Repeated measures ANOVA and Bayesian ANOVA inferential statistics for sex effects and interactions on social preference raw exploration and d2 scores.

| Analysis | Effect | *F* | *p* | $n^2_p$ | $BF_{exclusion}$ |
|---|---|---|---|---|---|
| Raw exploration time (s) | Item main effect | (1, 50) = 76.012 | < 0.001 | 0.603 | 0.000 |
| | Genotype main effect | (1, 50) = 0.014 | 0.907 | 0.000279 | 7.986 |
| | Sex main effect | (1, 50) = 0.108 | 0.744 | 0.002 | 6.845 |
| | Genotype × item | (1,50) = 0.479 | 0.492 | 0.009 | 7.678 |
| | Genotype × sex | (1, 50) = 1.737 | 0.193 | 0.034 | 9.851 |
| | Item × sex | (1, 50) = 0.391 | 0.535 | 0.008 | 3.923 |
| | Item × genotype × sex | (1, 50) = 0.003 | 0.957 | 0.0000577 | 46.918 |
| Discrimination ratio | Genotype main effect | (1, 50) = 0.850 | 0.361 | 0.017 | 4.973 |
| | Sex main effect | (1, 50) = 0.894 | 0.349 | 0.018 | 2.451 |
| | Genotype × sex | (1, 50) = 0.337 | 0.564 | 0.007 | 8.268 |

### S2.4 PCP-induced locomotion

In the pre-injection habituation period females habituated faster than males (significant time bin × sex interaction $F((2, 98) = 3.903)$, $p = 0.023$, $n^2_p = 0.074$) and females travelled less distance overall than males (sex main effect: $F((1, 49) = 8.302)$, $p = 0.006$, $n^2_p = 0.145$) however sex × genotype and sex × genotype × time bin interactions were non-significant as shown in Table S5. There were no sex effects in response to 5 mg/kg PCP with all sex effects and interactions non-significant as shown in Table S6.

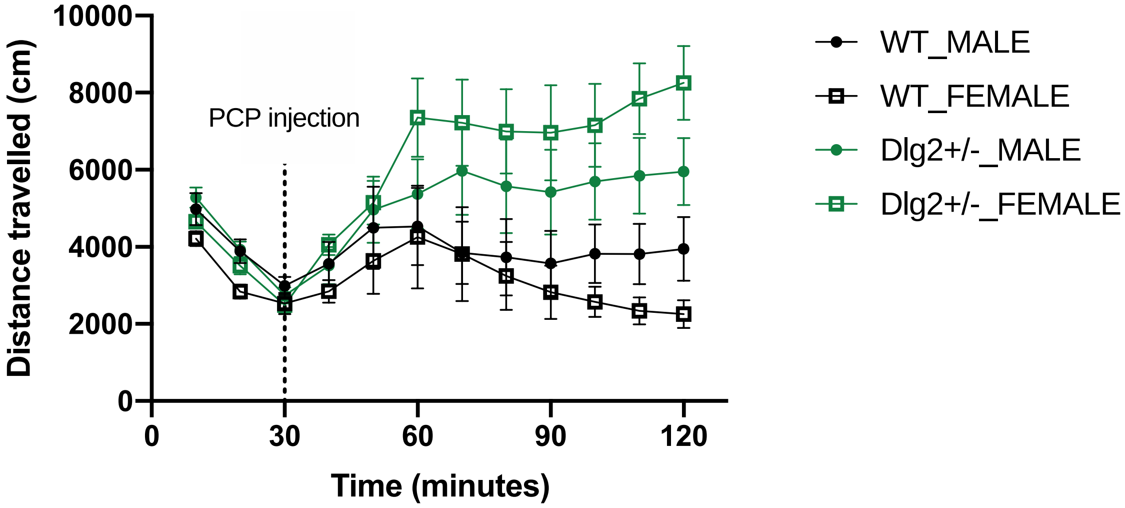


**Figure S6: Locomotor activity in response to PCP injection in male and female *Dlg2*$^{+/-}$ and wild-type rats.** Mean ± SEM distance travelled is plotted in 10-minute bins. The dotted line at 30 minutes denotes when the PCP injection occurred.

Table S5: Sex effects and interactions from repeated measures ANOVA and Bayesian repeated measures ANOVA of distance moved over the 30-minute pre-injection habituation period.

| Effect | *F* | *p* | $n^2_p$ | $BF_{exclusion}$ |
|---|---|---|---|---|
| Time bin main effect | (2, 98) = 227.098 | < 0.001 | 0.823 | < 0.001 |
| Sex main effect | (1, 49) = 8.302 | 0.006 | 0.145 | 0.064 |
| Genotype main effect | (1, 49) = 0.452 | 0.505 | 0.009 | 2.165 |
| Genotype × sex | (2, 98) = 1.077 | 0.304 | 0.022 | 1.596 |
| Time bin × sex | (2, 98) = 3.903 | 0.023 | 0.074 | 0.153 |
| Time bin × genotype | (2, 98) = 2.516 | 0.086 | 0.049 | 1.243 |
| Time bin × sex × genotype | (2, 98) = 0.298 | 0.743 | 0.006 | 4.156 |

Table S6: Sex effects and interactions from repeated measures ANOVA and Bayesian repeated measures ANOVA of distance moved over the 90-minute post-injection period.

| Effect | *F* | *p* | $n^2_p$ | $BF_{exclusion}$ |
|---|---|---|---|---|
| Time bin main effect | (2.829, 141.460) = 4.044 | 0.010 | 0.075 | < 0.001 |
| Sex main effect | (1, 50) = 0.183 | 0.670 | 0.004 | 3.328 |
| Genotype main effect | (1, 50) = 9.873 | 0.003 | 0.165 | < 0.001 |
| Genotype × sex | (2.829, 141.460) = 2.124 | 0.151 | 0.041 | 1.691 |
| Time bin × sex | (2.829, 141.460) = 0.714 | 0.537 | 0.014 | 57.096 |
| Time bin × genotype | (2.829, 141.460) = 5.125 | 0.003 | 0.093 | < 0.001 |
| Time bin × sex × genotype | (2.829, 141.460) = 0.786 | 0.497 | 0.015 | 305.148 |
